# ClpS directs degradation of primary N-end rule substrates in *Mycolicibacterium smegmatis*

**DOI:** 10.1101/2024.06.12.598358

**Authors:** Christopher J. Presloid, Jialiu Jiang, Pratistha Kandel, Henry R. Anderson, Patrick C. Beardslee, Thomas M. Swayne, Karl R. Schmitz

## Abstract

Drug-resistant tuberculosis infections are a major threat to global public health. The essential mycobacterial ClpC1P1P2 protease has received attention as a prospective target for novel antibacterial therapeutics. However, efforts to probe its function in cells are constrained by our limited knowledge of its physiological proteolytic repertoire. Here, we interrogate the role of mycobacterial ClpS in directing N-end rule proteolysis by ClpC1P1P2 in *Mycolicibacterium smegmatis*. Binding assays demonstrate that mycobacterial ClpS binds canonical primary N-degrons (Leu, Phe, Tyr, Trp) with moderate affinity. N-degron binding restricts the conformational flexibility of a loop adjacent to the ClpS N-degron binding pocket and strengthens ClpS•ClpC1 binding affinity ∼30-fold, providing a mechanism for cells to prioritize N-end rule proteolysis when substrates are abundant. Proteolytic reporter assays in *M. smegmatis* confirm degradation of substrates bearing primary N-degrons, but suggest that secondary N-degrons are absence in mycobacteria. This work expands our understanding of the mycobacterial N-end rule pathway and identifies ClpS as a critical component for substrate specificity, providing insights that may support the development of improved Clp protease inhibitors.

## INTRODUCTION

*Mycobacterium tuberculosis* has plagued civilization since antiquity and still causes greater global death than any other single infectious disease [1]. While most tuberculosis infections are treatable with existing drugs, over 400,000 new cases occur annually that exhibit drug-resistance, posing an ongoing challenge to global public health. New anti-tuberculosis therapeutics are required to address these drug-resistance infections. Mycobacterial Clp proteases, which carry out the regulated destruction of native cytosolic proteins, have received extensive attention as prospective antibacterial targets [2, 3]. These proteolytic machines consist of a ring-shaped unfoldase (ClpC1 or ClpX) that selects protein substrates, unravels them using energy from ATP hydrolysis, and passes them into an associated barrel-shaped hetero-oligomeric peptidase (ClpP1P2) for destruction [4–8]. All components of these enzymes are strictly essential in mycobacteria [7, 9–12], and several compound classes have been identified that kill *M. tuberculosis* in culture and in models of infection by dysregulating Clp proteases [2, 9, 10, 13–19]. Efforts to translate some lead compounds into clinically viable therapeutics are complicated by the risk of off-target activity towards the homologous mitochondrial CLPXP protease [20, 21]. However, humans lack an ortholog of the mycobacterial ClpC1 unfoldase, making this particular protease component an especially attractive antibacterial target. In spite of the essentiality of ClpC1P1P2, few physiological substrates have been described [5, 22–26]. An expanded understanding of ClpC1P1P2’s proteolytic repertoire would support work to identify its physiological roles and screen for compounds that disrupt its essential functions.

In many organisms, ATP-dependent proteases participate in N-end rule pathways that link the proteolytic half-life of cytosolic proteins to the identity of their N-terminal amino acid [27–29]. N-end rule substrates are typically delivered to proteases by N-end recognin adaptors [30, 31]. In *Escherichia coli*, for example, the adaptor ClpS recognizes substrates presenting N-terminal Leu, Phe, Tyr, or Trp residues and transfers them to the ClpA unfoldase of the ClpAP protease [30, 32, 33]. Similar N-end rule pathways mediated by ClpS and ClpA occur throughout Pseudomonadota (synonym Proteobacteria) [34–37], while ClpS cooperates with the ClpC unfoldase in Cyanobacteria [38], chloroplasts [39], and apicoplasts [40–42]. The set of residues that are directly recognized by ClpS – termed primary N-degrons – varies somewhat among species [34, 35] and is determined by the selectivity of a binding pocket on ClpS for hydrophobic side chains [32, 34, 36, 37, 43–45]. The landscape of N-end rule proteolysis is further diversified in many species by amino acid transferases that prepend primary N-degrons in front of other residues, creating secondary N-degrons [28, 35].

Recent studies point toward the existence of an N-end rule pathway in mycobacteria. Mycobacteria possess an ortholog of ClpS [46, 47] which has been shown to interact with ClpC1 [25]. Moreover, ClpS•ClpC1P1P2 degrades a model substrate bearing an N-terminal Phe *in vitro* [25]. However, physiological N-end rule proteolysis has not yet been observed in mycobacteria, and the complement of primary and secondary N-degrons is not known. Here, we use biochemical and structural approaches to examine the binding properties of mycobacterial ClpS, and employ a fluorescent reporter system to test for N-end rule proteolysis in *Mycolicibacterium smegmatis*, a model system for *M. tuberculosis*. We find that mycobacterial ClpS binds the four canonical primary N-degrons with moderate affinity, and that engagement with ClpC1 allosterically strengthens N-degron binding, likely through conformational stabilization of a mobile ClpS loop. Reporter assays establish that substrates bearing primary N-degrons are proteolyzed in *M. smegmatis*, but also indicate that secondary degrons are absent, in contrast to the well-established paradigm for N-end rule proteolysis in proteobacteria. This work expands our understanding of the physiological roles played by the essential ClpC1P1P2 protease in mycobacteria.

## METHODS

### Sequence analysis

**–** Orthologous sequences were compiled using the HMMER search algorithm [48] using *M. tuberculosis* ClpS (Uniprot ID: P9WPC1), *E. coli* ClpS (Uniprot ID: P0A8Q6), *E. coli* ClpA (Uniprot ID: P0ABH9), *M. tuberculosis* ClpC1 (Uniprot ID: P9WPC9), or *Streptomyces coelicolor* ClpIa (Uniprot ID: O69936) as a search query. Sequences were aligned with Clustal Omega [49] and pruned such that no two sequences had greater than 90% sequence identity. Expected secondary structure was annotated based on structures of the *E. coli* ClpS and the ClpA NTD (PDB: 1MBU) [33], the *M. tuberculosis* ClpC1 NTD (PDB: 6PBA) [50], or an AlphaFold2 [51] model of *Streptomyces coelicolor* ClpIa (Uniprot: O69936) [52]. Sequence alignments were visualized in Jalview [53]. Sequence logos were constructed using WebLogo [54]. Sequence conservation was mapped onto the *M. smegmatis* ClpS structure using Consurf [55, 56].

### Plasmid and strain construction

–Full-length ClpS coding regions were amplified from *E. coli* DH10β (NEB) and *M. smegmatis* MC^2^155 (ATCC 700084) genomic DNA and ligated into a derivative of pET-21a (EMD Biosciences) in frame with an upstream sequence encoding 6xHis-MBP-SUMO, providing a readily cleavable tag with a large size differential from the target protein. For crystallization of *^Msm^*ClpS, a variant lacking the N-terminal extension (*^Msm^*ClpS^core^; codons 21-100) was cloned into the same plasmid context. A synthetic DNA construct (Twist Biosciences) encoding full-length *M. tuberculosis* ClpC1 with *E. coli* codon optimization was cloned in frame with an N-terminal 7xHis-SUMO tag into a pET-22b derived vector.

Mycobacterial pTH shuttle plasmids were constructed by Gibson Assembly [57] from synthetic DNA fragments (Twist Biosciences) encoding a ColE1 *E. coli* origin of replication [58] from pET-22b, a pAL5000 mycobacterial origin of replication and hygromycin resistance cassette from pUV15tetORm [59], a TetR tetracycline expression cassette driven by the P*imyc* promoter [60], and a SUMO-X-mCherry^myc^ cassette driven by the tetracycline-regulated P*myc1-tet^On^* promoter (**Fig. S4**) [61]. A parallel set of pTHU vectors additionally incorporated an expression cassette for *Chaetomium thermophilum* Ulp1 SUMO protease [62] driven by the constitutive *M. tuberculosis Rv0005*p promoter [63]. The junction between SUMO and mCherry encoded the 12-residue sequence XLRVQSGTASGT [25, 30]; the initial “X” position was varied to each of the 20 amino acids. Integrating mycobacterial pTKI plasmids comprised ColE1, TetR, L5 integrase, and KanR components from PLJR962 [64], and a GFP-X-SUMO-mCherry^myc^ cassette driven by an optimized Ptet-opt promoter [64]; pTKIU plasmids additionally included the *Rv0005*p-Ulp1 cassette. All plasmid sequences were verified by Sanger (Genewiz) or nanopore sequencing (Plasmidsaurus or Genewiz).

### Protein purification

**–** Expression constructs were transformed into *E. coli* strain ER2566 (NEB), cultured to OD_600_ ≈ 0.6, and overexpression was induced by the addition of 0.5 mM isopropyl β-D-1-thiogalactopyranoside. *^Eco^*ClpS, *^Msm^*ClpS, and *^Msm^*ClpS^core^ constructs were expressed at 25°C for 5 h; *^Mtb^*ClpC1 was expressed overnight at 18°C. Cultures were centrifuged at 4000 rcf at 4°C for 30 min. The supernatant was discarded and the pellet from 1 L culture was re-suspended in 25 mL of Lysis Buffer (25 mM HEPES, 500 mM NaCl, 10 mM imidazole, 10% glycerol, 1mM phenylmethylsulfonyl fluoride, pH 7.5). Cells were lysed by sonication. His-tagged proteins were purified from clarified lysate by Ni-NTA (Marvelgent Biosciences). Yeast SUMO protease Ulp1 was used to remove the 6xHis-MBP-SUMO or 7xHis-SUMO tag at a ratio of ∼1:50 Ulp1 to target protein by overnight incubation at 4°C [65]. Cleaved protein was further purified by anion exchange (Source Q, Cytiva) and size exclusion chromatography (Superdex 75, Cytiva). *^Msm^*ClpS variants were stored in CPD (25 mM HEPES, 200 mM KCl, 10 mM MgCl_2_, 0.1 mM EDTA, pH 7.0); *^Eco^*ClpS was stored in APD (50 mM HEPES, 300 mM NaCl, 20 mM MgCl_2_, 10 % glycerol, 0.1 EDTA). Concentrations were determined by A_280_ readings on a NanoDrop1000 (Thermo Scientific). Reported concentrations of *^Mtb^*ClpC1 are hexamer equivalents.

### Proteolysis assays in M. smegmatis

**–** SUMO-X-mCherry^myc^ or GFP-SUMO-X-mCherry^myc^ reporter constructs were transformed by electroporation into *M. smegmatis* MC^2^155 at 2.5 kV in a 2 mm cuvette using a MicroPulser device (Bio-Rad). Liquid starter cultures were grown in 7H9 Middlebrook medium (7H9 Middlebrook base [HiMedia], 0.2% v/v glycerol [Fisher], 0.2% w/v glucose [TCI], 0.05% v/v Tween 80 [Fisher], 50 μg/mL hygromycin B [Goldbio] or 20 µg/mL kanamycin [Fisher]) with orbital shaking at 37°C. Starter cultures were sub-cultured into fresh media at a starting A_600_ of 0.05 in black clear bottomed Corning NBS 96-well plates. Reporter expression was induced by 50 ng/mL anhydrotetracycline (aTc). Fluorescence was monitored by 570 nm excitation and 640 nm emission for mCherry, or 468 nm excitation and 520 nm emission for GFP, and culture density was monitored by 600 nm absorbance in a Tecan Spark plate reader, with periodic shaking.

### Immunoblots

***–*** Liquid cultures of *M. smegmatis* harboring pTH or pTHU constructs were inoculated and grown for 48 h with shaking at 37°C. Cells were subcultured into 40 mL of fresh medium supplemented with 100 ng/mL aTc at a starting OD_600_ of 0.05. Samples were grown for 10 hours at 37°C, then harvested and pelleted at 1500 rcf for 15 min in 50 mL conical tubes. Supernatant was discarded and the pellet was resuspended in 3 mL Lysis Buffer. 650 μL of resuspended pellet slurry and 325 μL 0.1 mm zirconia/silica beads (BioSpec) were transferred to a microcentrifuge tube and lysed by bead beating in a Disruptor Genie (Scientific Industries) with 5 cycles of 1 min shaking followed by 30 s rest on ice. Lysed samples were centrifuged at 1500 rcf for 5 minutes to pellet beads and cellular debris, and the resulting supernatant was collected and stored at −20°C. 20 μL of thawed samples were run on SDS-PAGE gels and transferred to a polyvinylidene difluoride (PVDF) membrane. Blots were blocked at room temperature with 5% skim milk in TBS (20 mM Tris, 150 mM NaCl, pH 7.5) for 1 hour, then washed 3x in TBST (TBS with 0.1% Tween 20). Membranes were incubated with a 1:4000 dilution of mouse anti-cMyc antibody 9E10 (Invitrogen), followed by a 1:4000 dilution of goat anti-mouse IgG-HRP conjugate (Invitrogen), both in TBS with 3% milk. Blots were imaged using SuperSignal West Pico PLUS chemiluminescent substrate (Thermo) on a BioRad Chemidoc imager.

### Fluorescence anisotropy

**–** X-LFVQLASK^mal^ peptides bearing an N-terminal Leu, Phe, Tyr, Trp, or Ser and a maleimide-functionalized Lys were synthesized (Biomatik), along with a fluorescent CSGSK^TAMRA^ peptide bearing a C-terminal tetramethylrhodamine-functionalized Lys. Peptides were dissolved in 100 mM sodium phosphate pH 7.25, 10% v/v DMSO to 10 mM. Reactions consisting of 0.5 mM maleimide peptide and 0.25 mM TAMRA peptide in 100 mM sodium phosphate pH 7.25, 2.5% v/v DMSO were incubated for 1 h in the dark at room temperature, and quenched with 1 mM β-mercaptoethanol for 15 minutes. For anisotropy experiments, 0.1 µM of conjugated peptide was incubated with 0 – 100 µM ClpS in CPD (*^Msm^*ClpS) or APD (*^Eco^*ClpS) for 30 min, and fluorescence polarization readings were collected in a Tecan Spark plate reader. Data were fit by nonlinear regression to a quadratic single-site binding equation in Prism (Graphpad). Reported *K_D_* values are averages from 3 technical replicates.

### Microscale thermophoresis

– 7xHis-SUMO-*Mtb*ClpC1 was diluted to 2 µM in the presence of 20 µM ATPγS, RED-tris-NTA dye (NanoTemper), and 0.05% Tween 20 in CPD and incubated for 30 min at 20°C. Serial dilutions of *^Msm^*ClpS were prepared with or without inclusion of 200 µM TyrArg dipeptide in CPD+0.05% Tween 20, and were mixed 1:1 with *^Mtb^*ClpC1 solution. Thermophoretic changes were measure by the NanoTemper Monolith NT.115 at 40% LED and laser power at 25C. Data was fit to a hill equation and graphed in Prism (Graphpad).

### Crystallography

**–** Crystals of *^Msm^*ClpS^trunc^ were grown at 20°C by hanging drop vapor diffusion in drops consisting of 0.75 µL 7.5 mg/mL protein and 0.75 µL Crystallization Buffer (0.1 M HEPES, 12% PEG3350, 5 mM NiCl_2_, pH 7.5) over a 500 µL reservoir of Crystallization Buffer. Diamond shaped crystals (∼250 µm × 250 µm) were cryoprotected by soaking in Cryo Buffer (0.1 M HEPES, 40% PEG3350, 5 mM NiCl_2_, pH 7.5) for 1 h or overnight. For metal replacement soaks, MgCl_2_ or CoCl_2_ was substituted for NiCl_2_. For peptide soaks, 2.5 mM of peptide was included in the Cryo Buffer. X-ray diffraction datasets were collected at Rigaku MicroMax-007 HF or at ALS beamline 5.0.1. Data were processed with HKL-2000 [66]. The structure of *^Msm^*ClpS^trunc^ was solved by molecular replacement in Phaser [67] using the structure of *^Eco^*ClpS as a search model (PDB ID: 3O2O) [68]. Models were built in Coot [69] and refined in Phenix [70]. Crystallographic statistics are summarized in Table 1. Coordinates are available from the PDB (accession codes: 9AYN, 9AYO, 9AYP, 9AZ5, 9B06, 9B10, 9B1P).

**Table 1.**
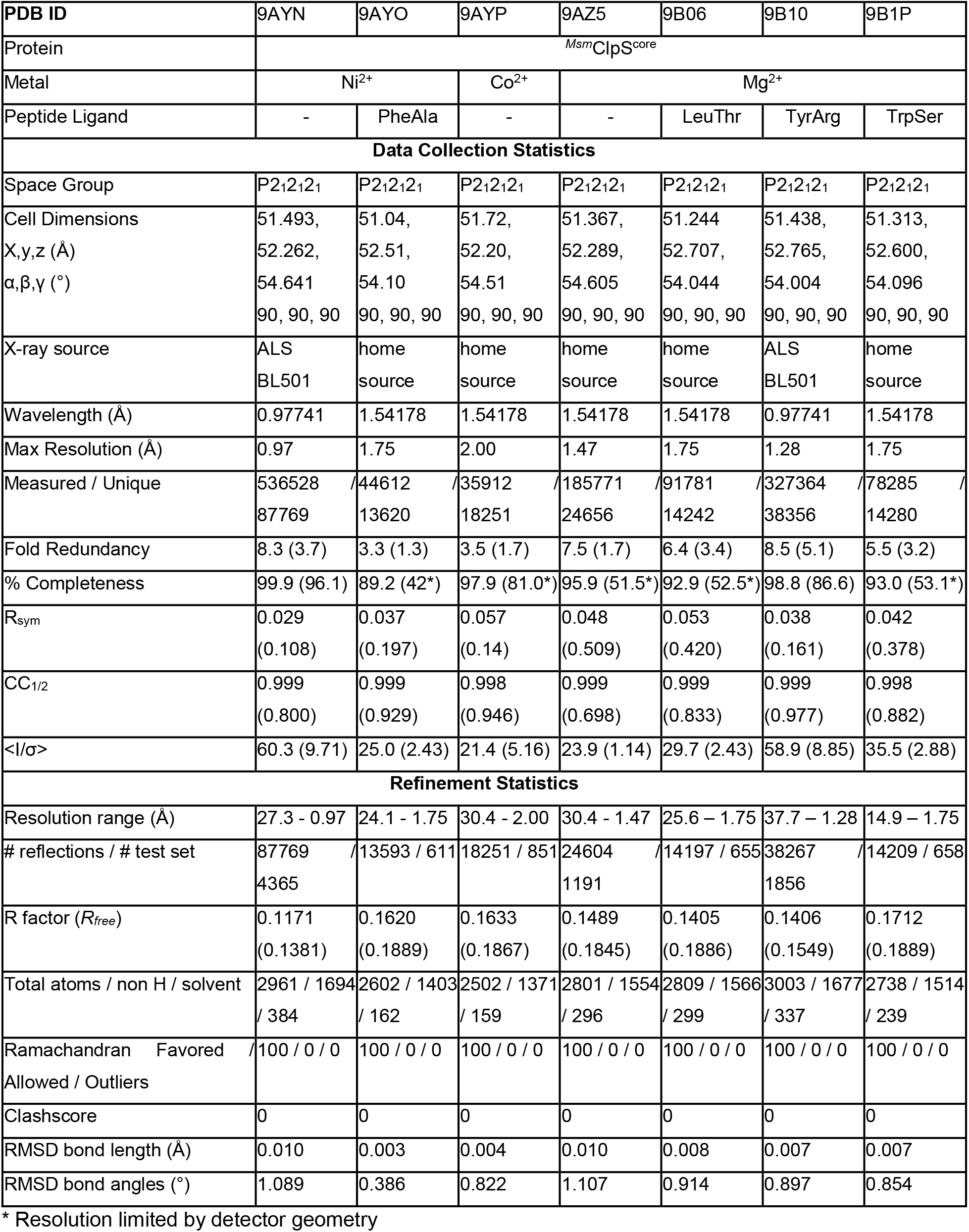
X-ray Crystal Structure Diffraction and Refinement Statistics.

### Flow cytometry

– *M. smegmatis* MC^2^155 were transformed with GFP-SUMO-X-mCherry^myc^ reporter constructs and grown in 7H9 Middlebrook medium (7H9 Middlebrook base [HiMedia], 0.2% v/v glycerol [Fisher], 0.2% w/v glucose [TCI], 0.05% v/v Tween 80 [Fisher], 20 μg/mL kanamycin [Fisher]) with orbital shaking at 37°C to saturation. Sub-cultures were prepared at 1:50 dilution into fresh media and grown at 37°C to mid-log phase. Subcultures were normalized to an OD_600_ of 0.05 with fresh media, and reporter expression was induced with 50 ng/ml anhydrotetracycline (aTc) at 37°C with shaking for 10 hours. 3×10^8^ cells were centrifuged for 5 min at 4000 g and resuspended in sterile Phosphate Buffered Saline (PBS) with 20 µg/ml kanamycin. mCherry fluorescence was monitored by excitation at 561 nm and GFP fluorescence by excitation at 488 nm in a BD FACSAria Fusion flow cytometer at the University of Delaware Flow Cytometry Core.

## RESULTS

### Sequence conservation of Actinomycetota ClpS orthologs

*-* Using *E. coli* ClpS (Uniprot: P0A8Q6) as a query, we identified ClpS orthologs within about half of the taxonomic orders within Actinomycetota (**Fig. 1A**). Notably, orthologs occur in *Mycobacterium tuberculosis* (Rv1331 in *M. tuberculosis* H37Rv) and *Mycolicibacterium smegmatis* (MSMEG_4910 in *M. smegmatis* mc^2^ 155), sharing 82% sequence identity. *E. coli* ClpS comprises two structural regions: a hypervariable ∼20 amino acid N-terminal segment required for substrate handoff to ClpA [68, 71, 72] followed by a conserved globular core [33, 44, 73]. Both regions are apparent in alignments of orthologs: the first 20 - 25 amino acids are present but poorly conserved, while the remaining ∼75 residues are conserved among Actinomycetota and are similar to *^Eco^*ClpS (∼30% identify to *^Msm^*ClpS), with the exception of a 6 amino acid gap near the C-terminus (**Fig. 1B**).

**Figure 1.**
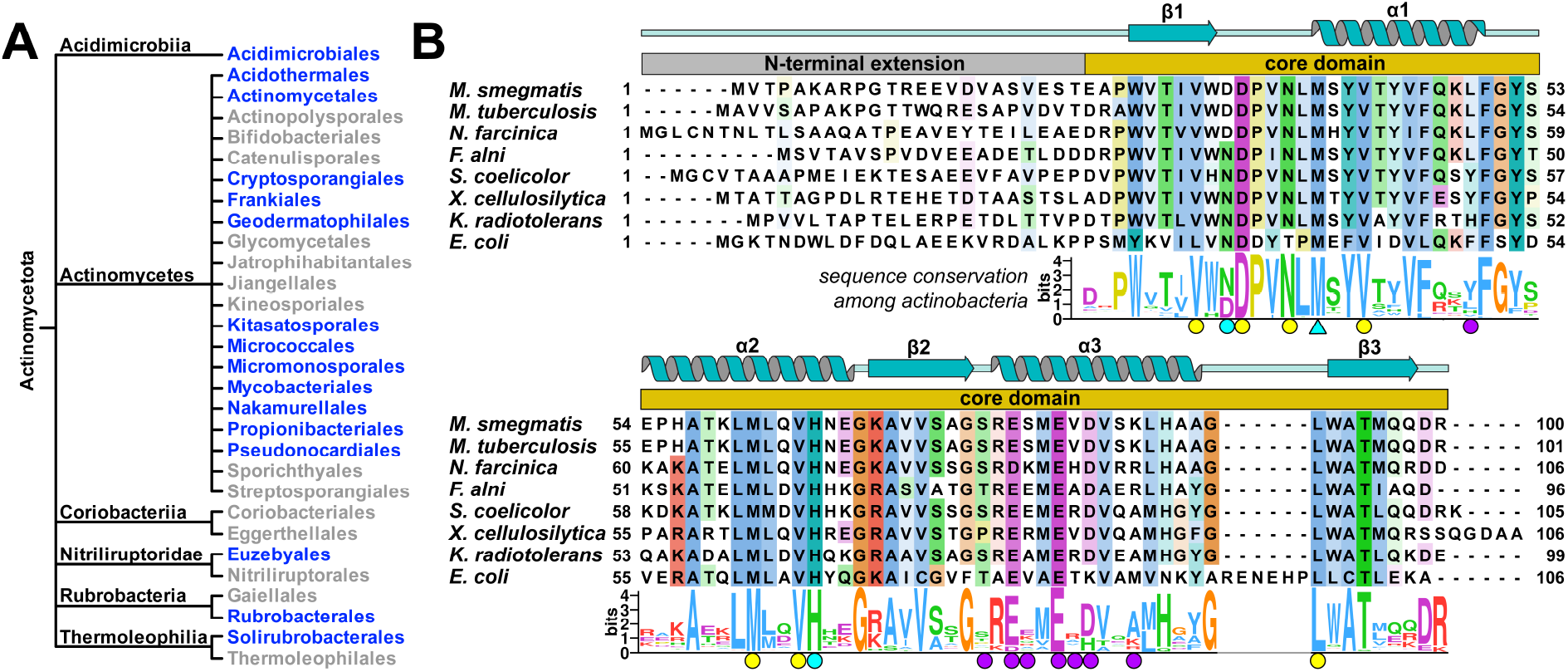
Distribution and sequence conservation of actinobacterial ClpS. **A**) ClpS orthologs are present (blue) in about half of the taxonomic orders within Actinomycetota. **B**) A sequence alignment shows conservation among ClpS orthologs from seven actinobacterial species and *E. coli*, with domain organization and expected secondary structure shown above, and a sequence logo below constructed from 230 actinobacterial orthologs. Residues that line the binding pocket (yellow circles), stabilize the N-degron amino-terminus (cyan circles), sterically block β-branched side chains (cyan triangle), or comprise the expected NTD-binding surface (purple circles) are noted.

The ClpS globular domain has two structurally defined interaction sites: *i*) a hydrophobic pocket that binds N-end rule side chains [32, 34, 36, 68] and *ii*) a surface that docks against the ClpA N-terminal domain (NTD) [33, 73]. We examined the conservation of ClpS residues associated with each interface, using *^Eco^*ClpS co-crystal structures as a reference. Positions in *^Eco^*ClpS that line the pocket (**Fig. 1B**, yellow circles) or contact the amino terminus of the N-degron (**Fig. 1B**, cyan circles) are strongly conserved, as is a key methionine that discriminates β-branched from unbranched side chains (Met40 in *E. coli*) [34, 36] (**Fig. 1B**, cyan triangle). These sequence features suggest that the orthologous pocket is similarly able to bind N-end rule motifs. By contrast, only piecemeal conservation was found among residues corresponding to the ClpA NTD-binding surface (**Fig. 1B**, purple circles). Two acidic amino acids that stabilize the interaction with ClpA in *E. coli* (equivalent to Glu78 and Glu81 in *^Msm^*ClpS) are strongly conserved, but flanking residues vary. The degree of conservation within this region is notably lower than observed among proteobacterial sequences (**Fig. S1A**).

Actinomycetota lack a direct ortholog of ClpA, but possess the architecturally related type-II AAA+ Clp unfoldases ClpC and ClpIa [5, 6, 52, 74]. The NTDs of both enzymes are structurally similar to that of ClpA [52]. A recent bacterial two-hybrid screen identified *M. tuberculosis* ClpS as a ClpC1 interaction partner, and demonstrated that ClpS delivers a model substrate bearing an N-terminal Phe (a primary N-degron in proteobacteria [27, 28, 34, 75]) to ClpC1P1P2 for degradation *in vitro* [25]. Given these findings, the low conservation of the putative NTD interaction surface is striking, especially because the ClpC NTD exhibits far greater conservation among Actinomycetota than does ClpA across Proteobacteria (**Fig. S1B,C**) [52]. This may indicate that the ClpS•NTD interface differs between clades. Alternatively, some ClpS orthologs may have evolved to interact with ClpIa, which occurs in only a subset of Actinomycetota species and possess greater sequence variability within its NTD (**Fig. 1D**).

### Crystal structures of M. smegmatis ClpS

*-* To assess its structural features, we determined the X-ray crystal structure of the *M. smegmatis* ClpS core domain (residues 21 - 100; *^Msm^*ClpS^core^), omitting the hypervariable N-terminal extension. Diffraction data were collected to a maximum resolution of 0.97 Å and the structure was solved by molecular replacement using *^Eco^*ClpS (PDB: 3O2O [68]) as a search model (**Table 1**). Two copies of *^Msm^*ClpS^core^ occupied the asymmetric unit (**Fig. 2A,B**), sharing similar overall conformations (backbone RMSD ∼0.4 Å) except for a small difference in the position of the β1-ɑ1 loop. Crystals required 5 mM NiCl_2_ to form. Once formed, crystals were stable for days in soaking conditions that replaced Ni^2+^ with Co^2+^ or Mg^2+^, but the addition of EDTA caused rapid cracking and dissolution. Several sites of electron density were observed near histidines and acidic side chains. An anomalous map generated from crystals soaked in Co^2+^ revealed a well-ordered Co^2+^ ion between chains in the asymmetric unit (coordinated by Asp99 on each chain) and additional Co^2+^ ions bridging symmetry related molecules (coordinated by Glu78 and Glu81 on one chain and His88 on the neighboring chain), thereby stabilizing crystal packing (**Fig. S2**). Little difference was observed among *^Msm^*ClpS^core^ structures determined from Ni^2+^, Co^2+^ or Mg^2+^ conditions (maximum RMSD ∼0.6 Å). (Notably, the Ni^2+^ ion bridging asymmetric units was present even after overnight incubation in Mg^2+^ conditions, indicating very stable coordination.)

*^Msm^*ClpS bears strong structural homology to proteobacterial ClpS (**Fig. 2C**), adopting an analogous ɑ/β fold with overall backbone RMSD between orthologs of ∼0.9 Å. The largest deviation is a shorter ɑ3-helix in *^Msm^*ClpS, corresponding to the 6-residue gap noted in the sequence alignment (**Fig. 1B**). Shortening of this helix eliminates contacts that buttress the β1-ɑ1 loop in *^Eco^*ClpS, likely accounting for the observed conformational variability between *^Msm^*ClpS chains. Analogous to *^Eco^*ClpS, *^Msm^*ClpS possesses a hydrophobic pocket bounded by β1, ɑ1, ɑ2, and the ɑ3-β3 loop (**Fig. 2D**). We mapped actionobacterial ClpS sequence conservation onto the surface of *^Msm^*ClpS using Consurf (**Fig. 2E**) [55, 56, 76]. As expected from the pattern of sequence conservation (**Fig. 1B**), surface residues surrounding the N-degron binding pocket are strongly conserved. Some of these define the shape of the pocket and potentially stabilize N-degron binding. Others (eg, Tyr41 and Trp32) are too distant to directly interact with a docked N-terminal residue. Visualization of surface electrostatics revealed a cluster of negative charge surrounding the pocket (**Fig. 2F**), which may help attract and orient positively charged termini of interacting N-degrons. Together, these observations indicate that the pocket is conserved across Actinomycetota. We also examined the region that corresponds to the ClpA NTD-interacting surface of *^Eco^*ClpS, based on co-crystal structures of *^Eco^*ClpS and the NTD of *^Eco^*ClpA (PDB: 1MG9, 1R6O) and found only partial conservation (**Fig. 2E**, circled).

**Figure 2.**
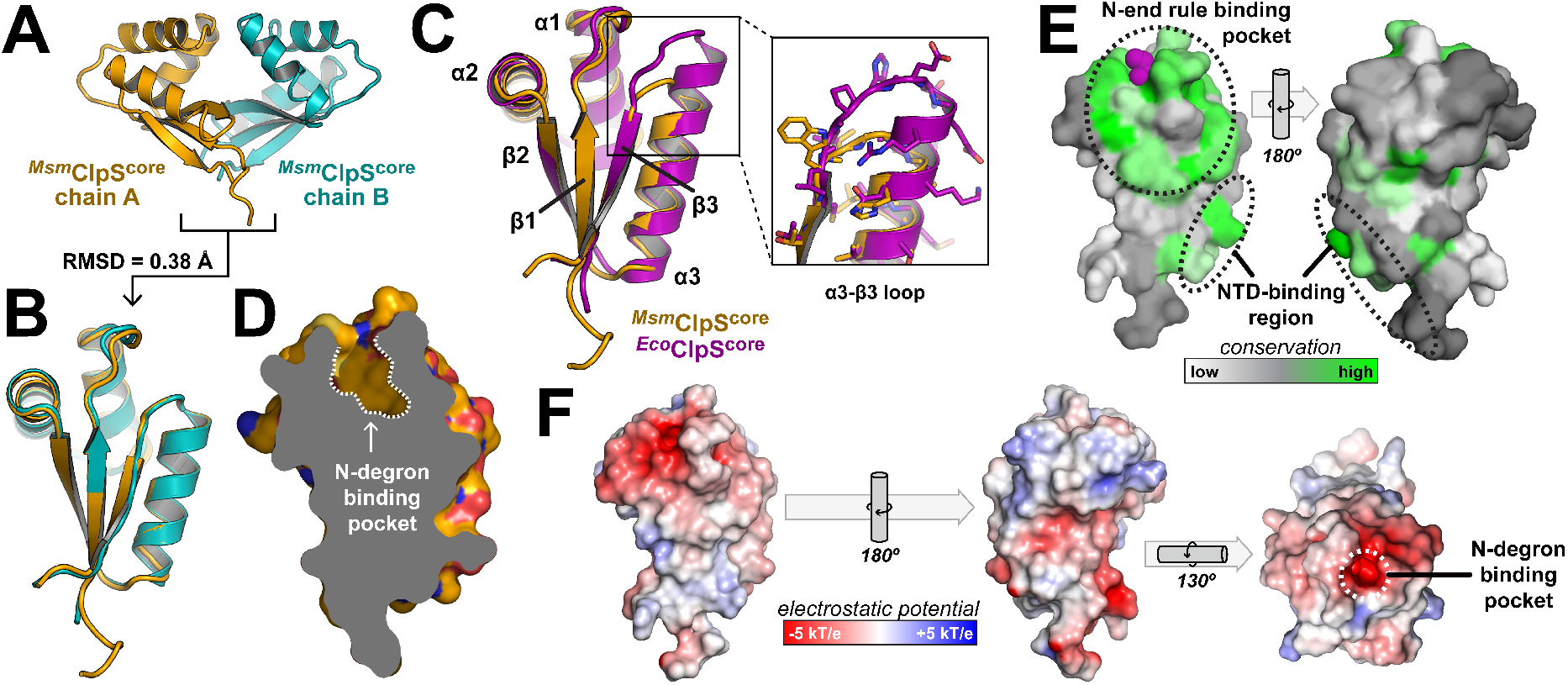
Crystal structure of *M. smegmatis* ClpS. **A**) Two copies of *^Msm^*ClpS^core^ occurred in the asymmetric unit, **B**) with low backbone deviation between chains. **C**) *M. smegmatis* ClpS^core^ (purple) adopts a similar fold to *E. coli* ClpS^core^ (gold) with the exception of a shorter α3 helix and α3-β3 loop. **D**) A cutaway view of *^Msm^*ClpS^core^ shows the N-degron binding pocket. **E**) The sequence conservation from an alignment of 230 ClpS orthologs was plotted on the surface of *^Msm^*ClpS^core^. **F**) The surface of *^Msm^*ClpS^core^ is shown colored by electrostatic potential, showing a region of negative charge around the N-degron binding pocket.

### Co-crystal structures with N-end rule peptides

– To examine details of N-end rule amino acid binding, crystal structures were determined of *^Msm^*ClpS^core^ in complex with dipeptides bearing canonical primary N-end rule residues. Dipeptides were introduced by soaking for 1 h in conditions containing either NiCl_2_ (PheAla) or MgCl_2_ (TrpSer, LeuThr), or overnight in MgCl_2_ (TyrArg). As in *E. coli* ClpS [32, 36, 44], the ɑ-carbon of the N-terminal residue rests at the entryway of the binding pocket, with the N-end rule side chain projecting within (**Fig. 3**, upper). Leu, Phe and Tyr dock in similar orientations through the ɣ-carbon, while the trajectory of Trp is rotated to accommodate its greater bulk. Peptide binding is stabilized by a salt bridge between the amino terminus and Asp33 (**Fig. 3**, lower). Additionally, His65 indirectly stabilizes peptides through a well-coordinated divalent metal ion that interacts simultaneously with the N-terminus, the carbonyl oxygen, and several bridging water molecules. When peptide soaks were performed in the presence of NiCl_2_, the stabilizing ion was best modeled as Ni^2+^. Interestingly, for peptides soaked in MgCl_2_, the identity of the ion depended on the length of the soak. In structures determined after soaking for ∼1 h, the ion remained Ni^2+^. Overnight incubation with MgCl_2_ allowed complete replacement of Ni^2+^ by Mg^2+^. Given the low abundance of free transition metal ions in cells, it is likely that this position is physiologically occupied by Mg^2+^, an abundant cation. However, it is interesting to note that mycobacterial ClpS has been implicated in the zinc-dependent proteolysis of the ribosomal hibernation regulator Mrf [77], so metal coordination at this site may be required for delivery of some substrates.

**Figure 3.**
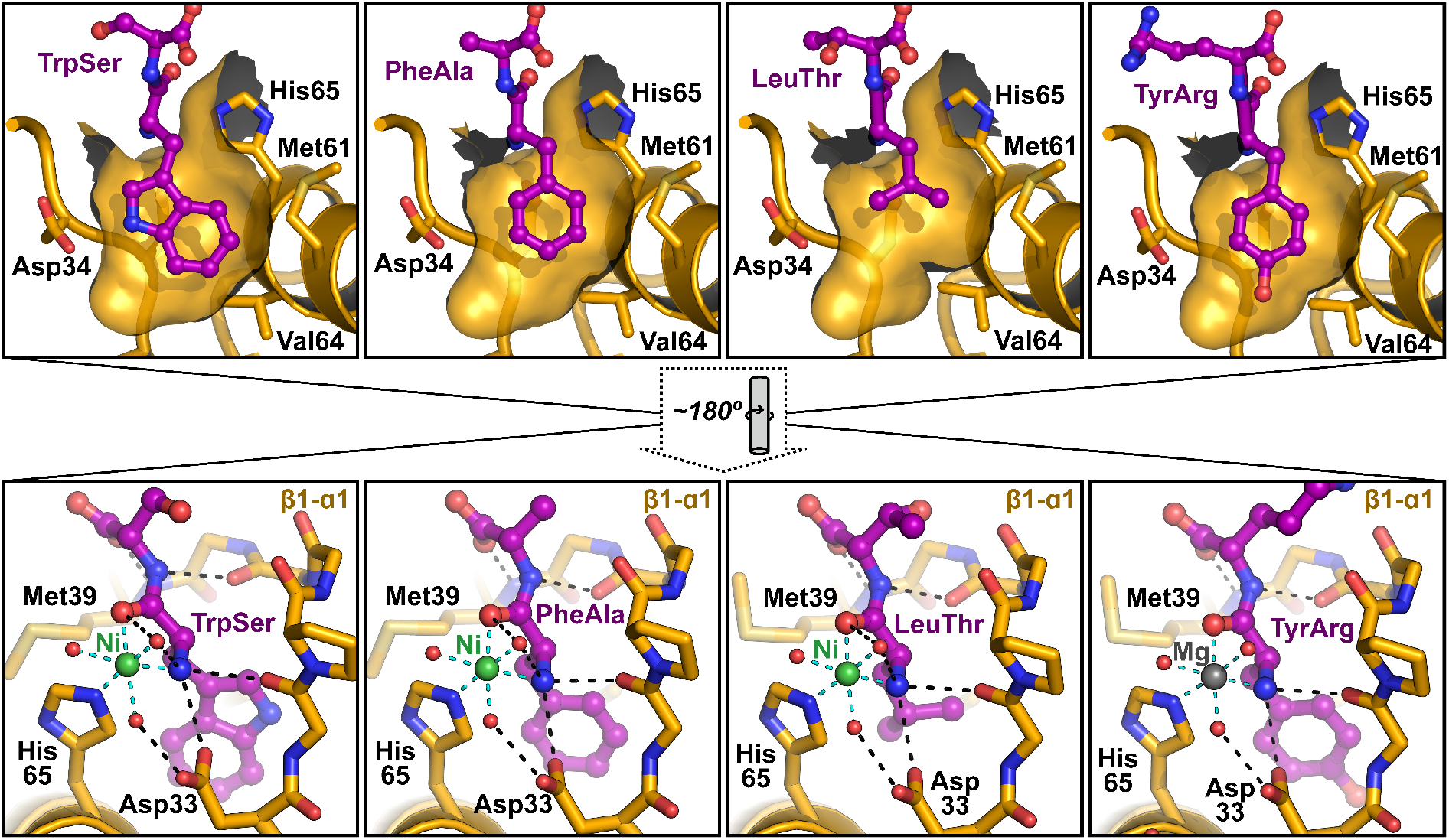
Crystal structures of N-end rule dipeptides bound to *M. smegmatis* ClpS. The indicated dipeptides (magenta) were co-crystallized with *M. smegmatis* ClpS^core^ (gold). Upper panels show peptides within the N-end rule binding pocket ligand, which is shown as a partial solvent-accessible surface. Lower panels show the upper entrance to the pocket, with a well-coordinated Ni^2+^ (green) or Mg^2+^ (gray) ion making bridging interactions between His65 and the peptide amino terminus. Notable residues are labeled, and polar contacts are shown as dashed lines.

The backbone of the β1-ɑ1 loop also contributes to peptide binding by forming hydrogen bonds with the dipeptide amino terminus and atoms of the second residue (**Fig. 3**, lower). This interaction in turn stabilizes a specific loop conformation. Whereas two distinct β1-ɑ1 trajectories occur in the ‘A’ and ‘B’ chains of peptide-free *^Msm^*ClpS^core^ structures, peptide binding induces a ∼2 Å shift in the ‘B’ loop to match the conformation of in ‘A’ (**Fig. 4A**). To systematically assess structural differences induced by peptide, we compared all ‘A’ and ‘B’ chains and found a consistent 0.3 - 0.4 Å difference in backbone RMSD between peptide-free and peptide-bound ‘B’ chains, but only minor differences in ‘A’ chains (**Fig. 4B**). We further constructed backbone atom distance matrices from representative peptide-free and peptide-bound ‘B’ chains (**Fig. 4C**), and plotted the absolute difference of inter-atom distances (**Fig. 4D**), which confirmed that the overall RMSD difference is due principally to peptide-induced shifts in the β1-ɑ1 loop. The fact that N-end rule peptide binding collapses a flexible loop into a single conformation suggests that binding incurs an entropic cost. There is thus an opportunity for interaction partners, such as the ClpC1 NTD, to modulate the strength of N-end rule degron binding (and vice versa) by influencing the β1-ɑ1 conformation.

**Figure 4.**
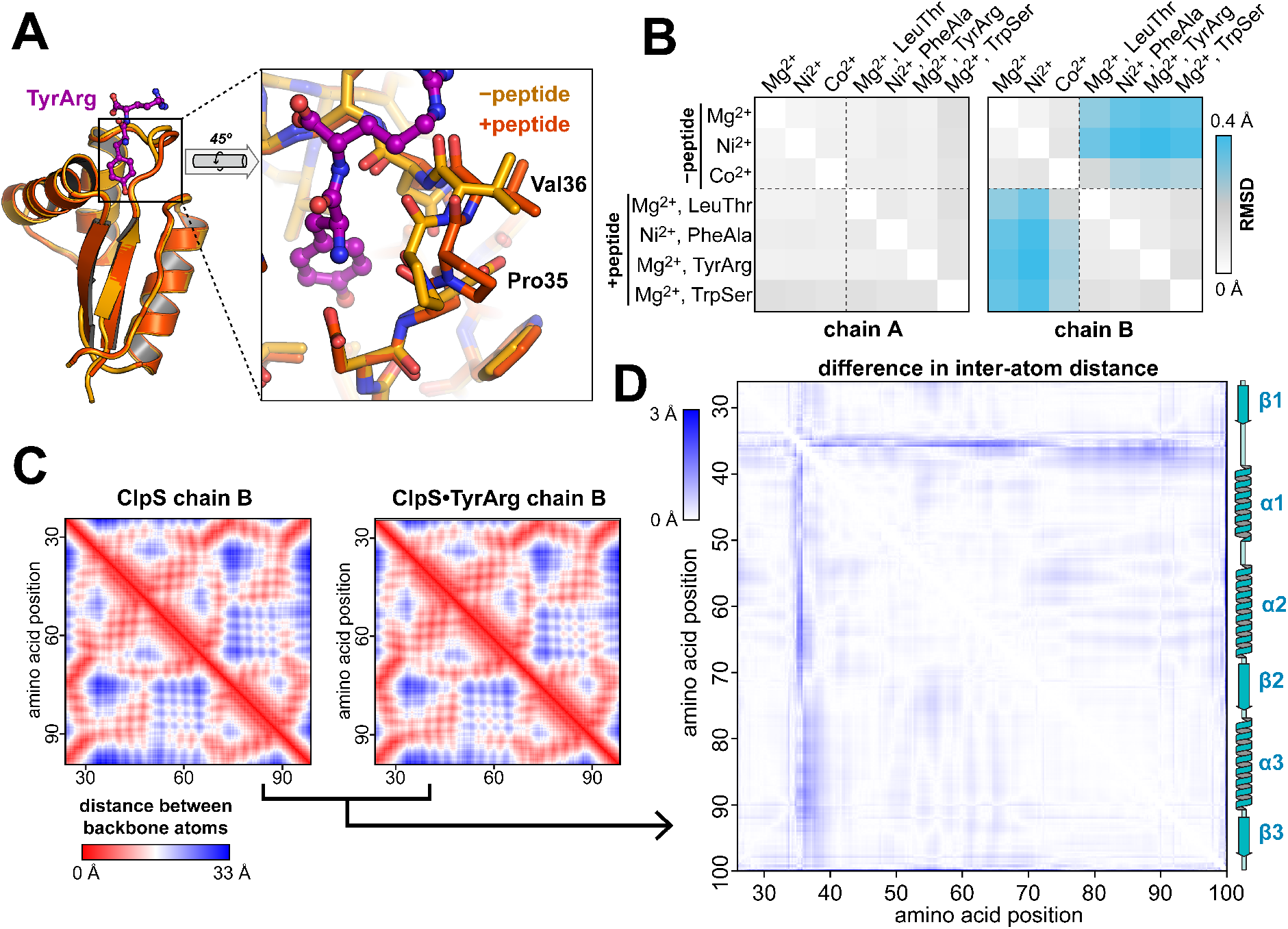
Peptide binding constrains the conformation of the α1/β1 loop. **A**) Superposition of chain ‘B’ from crystal structures determined in the presence of Mg^2+^ without (gold) and with (orange) bound TyrArg dipeptide. **B**) Pairwise root-mean square deviation of structures with and without bound peptides, determined separately for the ‘A’ and ‘B’ chains in the asymmetric unit. Increased RMSD is observed between peptide-bound and peptide-free ‘B’ chains. **C**) Heatmap illustrating intra-chain distances of main chain atoms from structures of ClpS in the presence of Mg^2+^ without (left) and with (right) TyrArg dipeptide. **D**) Heatmap of the absolute value of the differences in inter-atom distances between datasets in panel **C**. The largest differences are in the β1/α2 loop.

### Mycobacterial ClpS binds canonical primary N-end rule peptides

*-* To confirm that *^Msm^*ClpS binds N-end rule degrons, we measured fluorescence anisotropy of TAMRA-labeled peptides bearing varying N-terminal residues in the absence or presence of purified *^Msm^*ClpS (**Fig. 5A**; **Table 2**). No binding was observed for a control peptide with an N-terminal Ser. Peptides with canonical N-end rule amino acids in the leading position (Leu, Phe, Tyr, or Trp) showed increased anisotropy with increasing *^Msm^*ClpS, with *K_D_* ∼10 μM for aromatic N-terminal residues and 42 μM for a Leu-peptide. Notably, *^Msm^*ClpS dissociation constant were ≥ 10-fold larger than measured for *^Eco^*ClpS binding to the same peptides (**Fig. S3**; **Table 2**). These results confirm that *^Msm^*ClpS can bind canonical N-end rule degrons, in agreement with studies on *M. tuberculosis* ClpS [25, 47], however the relatively weak interaction may not be sufficient to support robust degradation in cells. Sequence determinants beyond the leading amino acid may strengthen binding for some N-terminal sequences. Alternatively, given the expected entropic cost of constraining the β1-ɑ1 loop (**Fig. 4**), we questioned whether N-degron binding to ClpS may be tighter in the presence of ClpC1. We tested this indirectly by measuring the interaction between ClpS and ClpC1 (bound to the slowly hydrolysable nucleotide ATPγS) in the absence and presence of an excess of TyrArg dipeptide, and found that TyrArg increased ClpS•ClpC1 binding affinity by nearly 30-fold (**Fig. 5B**). (A similar increase in affinity for N-degrons occurs when *E. coli* ClpS interacts with ClpA [68].) These results imply either thermodynamic linkage between ClpC1 binding and N-end rule substrate binding to ClpS, or that bound N-degron makes additional stabilizing contacts with part of ClpC1. Either model provides a mechanism by which accumulation of N-end rule substrates may prioritize ClpS-mediated proteolysis.

**Figure 5.**
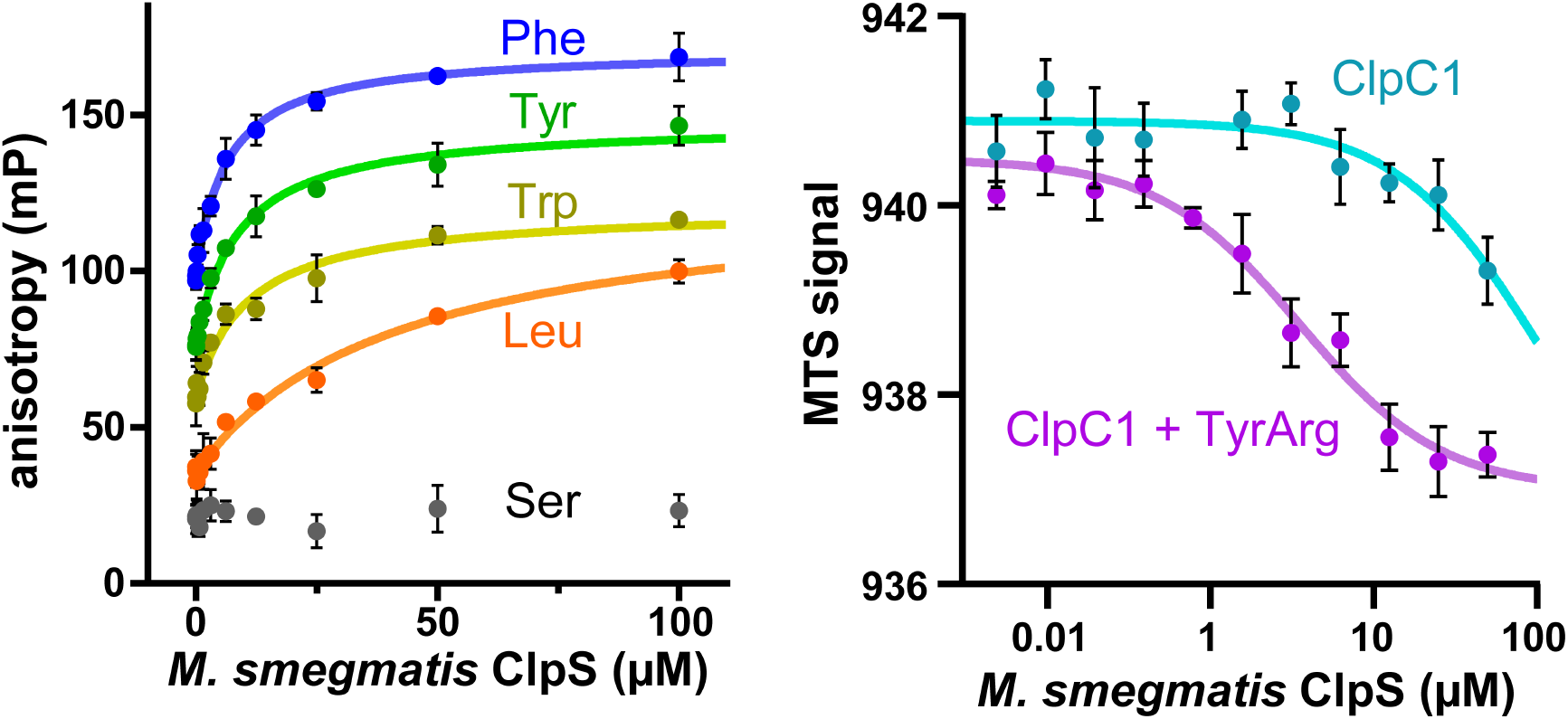
N-degrons and ClpC1 synergistically bind to ClpS. **A**) Fluorescence anisotropy of 100 nM TAMRA-labeled peptides bearing the indicated N-terminal amino acid was monitored over increasing concentrations of *M. smegmatis* ClpS. Data were fit to a single site binding equation. ClpS bound canonical N-degron peptides harboring Leu with *K_D_* = 42 ± 9 µM, Tyr with *K_D_* = 8.8 ± 1 µM, Trp with *K_D_* = 11 ± 2 µM, Phe with *K_D_* = 6.5 ± 1 µM. No binding was observed to a peptide with N-term Ser. **B**) Binding between *M. smegmatis* ClpS and 1 µM ClpC1•ATPγS, in the presence or absence of 200 µM TyrArg dipeptide, was monitored by microscale thermophoresis. Data were fit to a non-cooperative single site binding equation. ClpS bound ClpC1 with *K_app_* = 100 ± 160 µM in the absence of TyrArg, and *K_app_* = 3.5 ± 0.7 µM in the presence of TyrArg.

**Table 2.** Peptide binding constants vs ClpS.

|  | *Msm*ClpS |  | *Eco*ClpS |  |
| --- | --- | --- | --- | --- |
| | $K_D$ (μM) | $R^2$ | $K_D$ (μM) | $R^2$ |
| Leu-peptide | $42 \pm 8.8$ | 0.956 | $0.88 \pm 0.07$ | 0.990 |
| Phe-peptide | $6.5 \pm 0.9$ | 0.972 | $1.0 \pm 0.08$ | 0.991 |
| Tyr-peptide | $8.8 \pm 1.3$ | 0.969 | $2.5 \pm 0.2$ | 0.991 |
| Trp-peptide | $11 \pm 2$ | 0.945 | $0.95 \pm 0.08$ | 0.987 |
| Ser-peptide | n.b. | - | n.b. | - |

n.b. – no binding

### Primary N-end rule substrates are proteolyzed in M. smegmatis

– Our structural and biochemical data demonstrate that mycobacterial ClpS can directly bind the four canonical primary N-end rule amino acids. Proteobacterial N-end rule pathways additionally process proteins that present secondary N-end rule residues (eg, Arg and Lys in *E. coli*) which are converted by amino acid transferases into primary degrons through the addition of a Leu or Phe to the N-terminus [28, 35]. While orthologs of *E. coli* Leu/Phe-tRNA-protein transferase (aat) [78, 79] are present in some Actinomycetota species, we found no identifiable homologs of aat or the *V. vulnificus* amino acid transferase bpt [35] in Mycobacteriales, calling into question whether secondary N-degrons exist in mycobacteria.

To empirically test for primary and secondary N-degrons in *M. smegmatis*, we constructed a panel of fluorescent reporter substrates with varying N-terminal residues shielded by a removable tag (**Fig. S4**). The sequence X-RSKGEELVTGT [30] was cloned in front of an mCherry construct carrying a C-terminal myc tag, with the “X” position varied to each of the 20 amino acids. The “X” residue was shielded by an upstream *S. cerevisiae* SUMO domain [65], similar to the ubiquitin shield used in pioneering *E. coli* N-end rule studies by Varshavsky and colleagues [28]. Plasmids encoding SUMO-X-mCherry^myc^ were constructed with and without a separate expression cassette encoding the SUMO protease Ulp1 [62, 65]. Immunoblots confirmed that co-expression of Ulp1 led to removal of the shielding SUMO tag (**Fig. 6A, Fig. S5**) in all constructs except SUMO-Pro-mCherry^myc^, presumably due to the inability of Ulp1 to cleave when Pro occupies the P1 position [65]. Use of the SUMO shield avoids ambiguity surrounding removal of the initiator Met by methionine aminopeptidase [80], and ensures presentation of the intended N-terminal residue.

**Figure 6.**
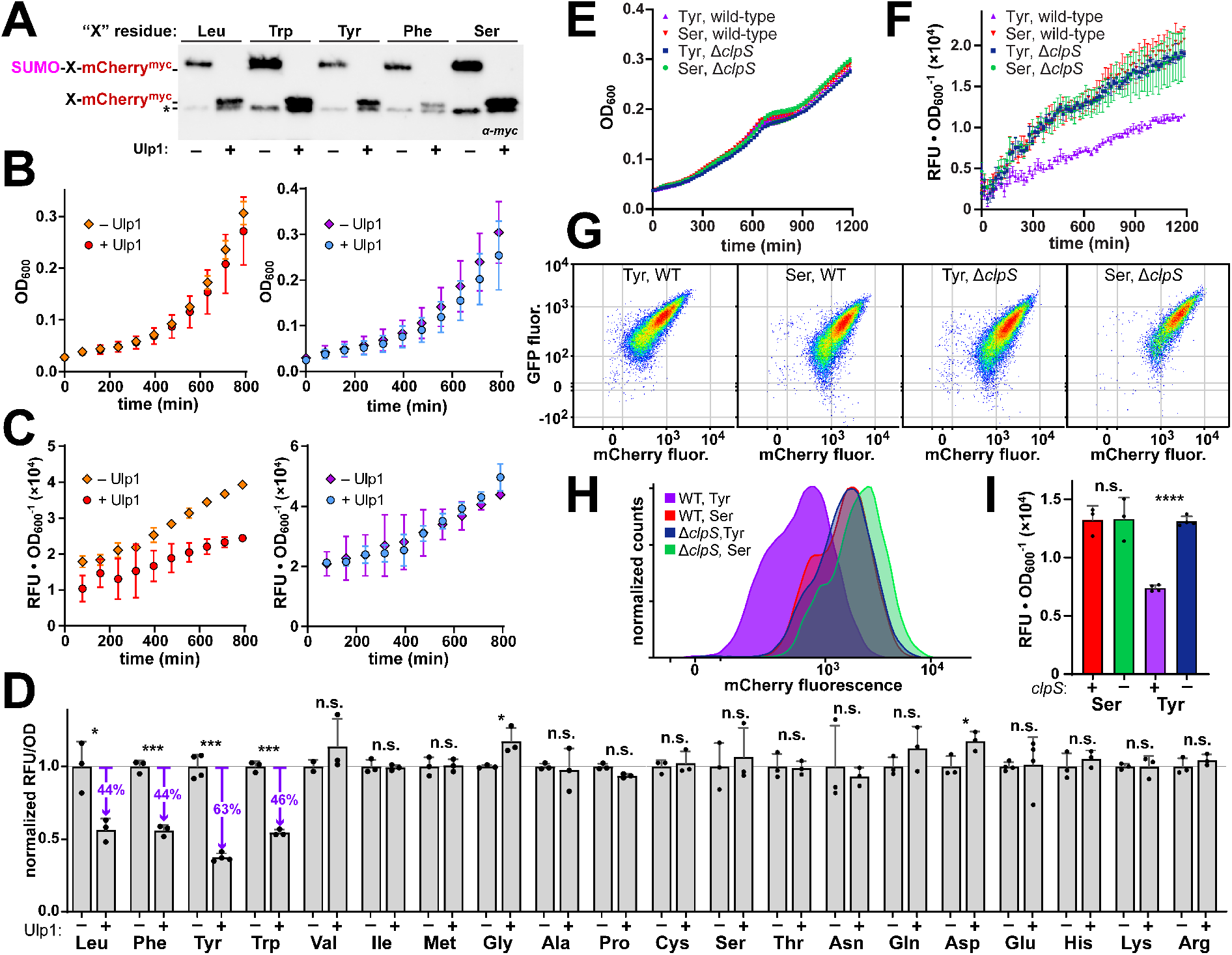
Physiological ClpS-dependent N-end rule proteolysis occurs in *M. smegmatis*. **A**) Immunoblots of lysates from cells expressing the indicated SUMO-X-mCherry^myc^ constructs, with or without co-expressed Ulp1, were probed with an α-myc antibody. Constructs migrates at as smaller species in the presence of Ulp1, consistent with removal of the shielding SUMO. The asterisk indicates a truncation product that incorporates most of mCherry. See also Fig. S5. **B**) Growth curve of cells expressing SUMO-Tyr-mCherry^myc^ (left) or SUMO-Ser-mCherry^myc^ (right), with or without Ulp1 co-expression, shows similar growth. **C**) Plots of mCherry fluorescence normalized by OD of samples in panel **B** shows lower fluorescence of SUMO-Tyr-mCherry^myc^ when Ulp1 is co-expressed. **D**) The normalized mCherry fluorescence of cells expressing SUMO-X-mCherry^myc^ constructs with the indicated residue in the “X” position, with or without Ulp1 co-expression, after 10 h growth. Leu, Phe, Tyr and Trp constructs exhibit large statically significant reductions in fluorescence in the context of Ulp1 co-expression. **E**) Growth curves of wild-type and Δ*clpS* cells expressing GFP-SUMO-Tyr-mCherry^myc^ or GFP-SUMO-Ser-mCherry^myc^, with Ulp1 co-expression, shows similar growth. **F**) Plots of mCherry fluorescence normalized by OD of cultures in panel **F** shows lower fluorescence of GFP-SUMO-Tyr-mCherry^myc^ only in wild-type cells. **G**) Flow cytometry analysis of cultures in panel **F**, plotting GFP fluorescence against mCherry fluorescence, shows a shift to low mCherry fluorescence signal for wild-type cells expressing GFP-SUMO-Tyr-mCherry^myc^ (right) compared to all other samples, but similar GFP fluorescence levels. See also Fig. S6. **H**) Normalized histograms of mCherry fluorescence from panel **G** show that deletion of *clpS* causes cells expressing GFP-SUMO-Tyr-mCherry^myc^ to adopt fluorescence levels similar to non-proteolyzed Ser controls. **I**) The normalized mCherry fluorescence values of cells expressing GFP-SUMO-Tyr-mCherry^myc^ or GFP-SUMO-Ser-mCherry^myc^, with Ulp1 co-expression, after 10 h, show a statistically significant increase in Tyr fluorescence when *clpS* is absent. Error bars reflect SD from ≥ 3 biological replicates. p-values were calculated by unpaired two-tailed Student’s t-test. ns: p > 0.05; * p ≤ 0.05, *** p ≤ 0.001, **** p ≤ 0.0001

*M. smegmatis* cultures grew similarly regardless of the amino acid in the variable position or the presence/absence of Ulp1 (**Fig. 6B**). Comparing steady-state mCherry expression, we found that most constructs exhibited indistinguishable fluorescence levels in the presence and absence of Ulp1. However, constructs bearing N-terminal Leu, Phe, Tyr and Trp produced 44 - 64% lower fluorescence when Ulp1 was present, compared to its absence (**Fig. 6C,D**), consistent with N-end rule proteolysis of substrates bearing these canonical primary N-degrons. (Removal of SUMO resulted in small but statistically significant increases in signal for Gly and Asp constructs, which may reflect packing interactions that influence mCherry brightness.) Importantly, constructs bearing other N-terminal residues accumulated to similar levels in the presence and absence of Ulp1, suggesting that secondary N-degrons are absent, at least under these growth conditions.

To test whether N-end rule proteolysis requires ClpS, we used CRISPR [81, 82] to generate a *M. smegmatis* strain lacking *clpS*. We also modified the Tyr and Ser reporter constructs to incorporate a GFP fusion in front of SUMO (GFP-SUMO-X-mCherry^myc^) (**Fig. S4**), allowing us to follow GFP fluorescence as a readout for construct expression [45]. Deletion of *clpS* did not alter cell growth (**Fig. 6E**). Plate reader assays (**Fig. 6F,I**) and flow cytometry analysis (**Fig. 6G,H, Fig. S6**) demonstrated that Ser reporters produced similar fluorescence levels in wild-type and Δ*clpS* cells. By contrast, the strong reduction in signal observed in wild-type cells with the Tyr reporter was abolished in Δ*clpS* cells, confirming that N-end rule proteolysis requires ClpS.

## DISCUSSION

Our findings demonstrate that mycobacterial ClpS functions physiologically as an N-recognin for proteolysis of N-end rule substrates. As in *E. coli*, mycobacterial ClpS is non-essential [11, 12], and likely plays a dispensable role in protein quality control by eliminating the products of aberrant protein cleavage events, and also contributes to regulated proteolysis of specific targets, such as the ribosome remodeling factor Mrf [77]. Interestingly, the sequence conservation of actinobacterial ClpS is markedly lower than that of ClpC overall or the ClpC NTD specifically, suggesting that ClpS-dependent pathways are not a major driver of ClpC NTD conservation, and that the most conserved functions of ClpC involve other substrates or interaction partners. Conservation patterns within the NTD of ClpIa, a type-II Clp unfoldase paralog present in a subset of actinobacteria [52], raise the possibility that some ClpS orthologs interact with ClpIa (or yet other protease systems) instead of, or in addition to, ClpC1. This may account for the diversity in ClpS surface residues, and allow some actinobacterial species to uncouple dispensable N-end rule proteolysis from the growth-essential functions of ClpC1.

Our experiments reveal additive effects between binding of N-degron peptides and ClpC1 to ClpS, with N-degrons increasing the affinity of ClpS for ClpC1 by ∼30-fold. By comparing ClpS structures with and without bound peptides – generated from essentially identical crystallization conditions – we see evidence that N-degron binding stabilizes the conformationally flexible of the β1-ɑ1 loop. This provides a possible mechanism through which either binding partner may stabilize a higher affinity conformation of ClpS. A similar high-affinity ternary complex was observed with *E. coli* N-end rule components [68], although it is unclear if the underlying mechanism is the same. High-affinity binding of *E. coli* ClpS to ClpA involves engagement of the ClpS N-terminal extension, and is thought to additionally involve contacts between an engaged N-degron and elements of the ClpA D1-ring [68, 72, 83]. A prior study identified interactions between mycobacterial ClpS and the ClpC1 M-domain, which may similarly contribute to high-affinity ClpS•ClpC1 binding [74]. By stabilizing the ClpS•ClpC1P1P2 proteolytic complex, N-degrons may prioritize ClpS-dependent N-end rule proteolysis under conditions where such substrates are abundant, and allow dissociation of ClpS following N-degron clearance. Importantly, ClpS has been shown to suppress proteolysis of a non-N-end rule substrate by ClpC1P1P2 [25]. The ability to promote dissociation of ClpS from ClpC1P1P2 when unneeded may be particularly important for mycobacteria, given the essentiality of non-ClpS-dependent proteolytic pathways.

One major difference between the *M. smegmatis* N-end rule pathway defined here and the well-characterized N-end rule pathways of proteobacteria [35, 78] is the absence of secondary N-degrons, which is in accordance with the lack of mycobacterial aat and bpt amino acid transferase homologs. Aside from expanding the number of destabilizing N-terminal residues, amino acid transferases that create secondary N-degrons provide a potential point of regulation over proteobacterial protein half-life. It is possible that mycobacteria instead regulate N-degron exposure through the use of alternative methionine aminopeptidase (map) enzymes that selectively remove the deformylated initiator methionine from nascent polypeptides [84]. *E. coli* possesses a single map ortholog that efficiently removes the leading Met when the second residue is small (Ala, Cys, Gly, Pro, Ser) but leaves Met intact before other residues, [84–86] and thus avoids constitutively creating N-degrons. Mycobacteria typically possess multiple map paralogs, with two in *M. tuberculosis* H37Rv (mapA/Rv0734 and mapB/Rv2861c) [87] and four in *M. smegmatis* mc^2^155 (MSMEG_2587, MSMEG_5050, MSMEG_5683, and MSMEG_1485). If these enzymes have varied specificity for downstream residues, it may provide a mechanism by which mycobacteria can modulate N-degron exposure.

Our research adds to the relatively short list of physiological substrates known for ClpC1P1P2, and suggests that mycobacteria possess a simpler N-end rule landscape than observed in proteobacteria. Future work will be required to understand whether mycobacteria exert regulatory control over N-degron exposure, and to what extent N-degron proteolysis is prioritized over the various classes of other substrates that interact with the highly conserved ClpC1 NTD. Importantly, model substrates incorporating N-degrons may serve as useful reporters of ClpC1P1P2 activity for screening applications and mechanistic studies of ClpC1-targeting inhibitors.

## References

1. WHO: Global tuberculosis report 2023. In. Edited by Organization WH. Geneva: World Health Organization; 2023.

2. Lee H, Suh JW: Anti-tuberculosis lead molecules from natural products targeting Mycobacterium tuberculosis ClpC1. Journal of industrial microbiology & biotechnology 2016, 43(2-3):205–212.

3. Xu X, Zhang L, Yang T, Qiu Z, Bai L, Luo Y: Targeting caseinolytic protease P and its AAA1 chaperone for tuberculosis treatment. Drug discovery today 2023, 28(3):103508.

4. Leodolter J, Warweg J, Weber-Ban E: The Mycobacterium tuberculosis ClpP1P2 Protease Interacts Asymmetrically with Its ATPase Partners ClpX and ClpC1. PloS one 2015, 10(5):e0125345.

5. Schmitz KR, Sauer RT: Substrate delivery by the AAA+ ClpX and ClpC1 unfoldases activates the mycobacterial ClpP1P2 peptidase. Molecular microbiology 2014, 93(4):617–628.

6. Akopian T, Kandror O, Raju RM, Unnikrishnan M, Rubin EJ, Goldberg AL: The active ClpP protease from M. tuberculosis is a complex composed of a heptameric ClpP1 and a ClpP2 ring. The EMBO journal 2012, 31(6):1529–1541.

7. Raju RM, Unnikrishnan M, Rubin DH, Krishnamoorthy V, Kandror O, Akopian TN, Goldberg AL, Rubin EJ: Mycobacterium tuberculosis ClpP1 and ClpP2 function together in protein degradation and are required for viability in vitro and during infection. PLoS Pathog 2012, 8(2):e1002511.

8. Benaroudj N, Raynal B, Miot M, Ortiz-Lombardia M: Assembly and proteolytic processing of mycobacterial ClpP1 and ClpP2. BMC Biochem 2011, 12:61.

9. Famulla K, Sass P, Malik I, Akopian T, Kandror O, Alber M, Hinzen B, Ruebsamen-Schaeff H, Kalscheuer R, Goldberg AL et al: Acyldepsipeptide antibiotics kill mycobacteria by preventing the physiological functions of the ClpP1P2 protease. Molecular microbiology 2016, 101(2):194–209.

10. Ollinger J, O’Malley T, Kesicki EA, Odingo J, Parish T: Validation of the essential ClpP protease in Mycobacterium tuberculosis as a novel drug target. J Bacteriol 2012, 194(3):663–668.

11. DeJesus MA, Gerrick ER, Xu W, Park SW, Long JE, Boutte CC, Rubin EJ, Schnappinger D, Ehrt S, Fortune SM et al: Comprehensive Essentiality Analysis of the Mycobacterium tuberculosis Genome via Saturating Transposon Mutagenesis. mBio 2017, 8(1).

12. Sassetti CM, Boyd DH, Rubin EJ: Genes required for mycobacterial growth defined by high density mutagenesis. Molecular microbiology 2003, 48(1):77–84.

13. Choules MP, Wolf NM, Lee H, Anderson JR, Grzelak EM, Wang Y, Ma R, Gao W, McAlpine JB, Jin YY et al: Rufomycin Targets ClpC1 Proteolysis in Mycobacterium tuberculosis and M. abscessus. Antimicrobial agents and chemotherapy 2019, 63(3).

14. Gao W, Kim JY, Anderson JR, Akopian T, Hong S, Jin YY, Kandror O, Kim JW, Lee IA, Lee SY et al: The cyclic peptide ecumicin targeting ClpC1 is active against Mycobacterium tuberculosis in vivo. Antimicrobial agents and chemotherapy 2015, 59(2):880–889.

15. Moreira W, Ngan GJ, Low JL, Poulsen A, Chia BC, Ang MJ, Yap A, Fulwood J, Lakshmanan U, Lim J et al: Target mechanism-based whole-cell screening identifies bortezomib as an inhibitor of caseinolytic protease in mycobacteria. mBio 2015, 6(3):e00253–00215.

16. Bürstner N, Roggo S, Ostermann N, Blank J, Delmas C, Freuler F, Gerhartz B, Hinniger A, Hoepfner D, Liechty B et al: Gift from Nature: Cyclomarin A Kills Mycobacteria and Malaria Parasites by Distinct Modes of Action. Chembiochem : a European journal of chemical biology 2015, 16(17):2433–2436.

17. Gavrish E, Sit CS, Cao S, Kandror O, Spoering A, Peoples A, Ling L, Fetterman A, Hughes D, Bissell A et al: Lassomycin, a ribosomally synthesized cyclic peptide, kills mycobacterium tuberculosis by targeting the ATP-dependent protease ClpC1P1P2. Chem Biol 2014, 21(4):509–518.

18. Schmitt EK, Riwanto M, Sambandamurthy V, Roggo S, Miault C, Zwingelstein C, Krastel P, Noble C, Beer D, Rao SP et al: The natural product cyclomarin kills Mycobacterium tuberculosis by targeting the ClpC1 subunit of the caseinolytic protease. Angewandte Chemie (International ed in English) 2011, 50(26):5889–5891.

19. Compton CL, Schmitz KR, Sauer RT, Sello JK: Antibacterial activity of and resistance to small molecule inhibitors of the ClpP peptidase. ACS chemical biology 2013, 8(12):2669–2677.

20. Wong KS, Mabanglo MF, Seraphim TV, Mollica A, Mao YQ, Rizzolo K, Leung E, Moutaoufik MT, Hoell L, Phanse S et al: Acyldepsipeptide Analogs Dysregulate Human Mitochondrial ClpP Protease Activity and Cause Apoptotic Cell Death. Cell chemical biology 2018, 25(8):1017–1030 e1019.

21. Cole A, Wang Z, Coyaud E, Voisin V, Gronda M, Jitkova Y, Mattson R, Hurren R, Babovic S, Maclean N et al: Inhibition of the Mitochondrial Protease ClpP as a Therapeutic Strategy for Human Acute Myeloid Leukemia. Cancer cell 2015, 27(6):864–876.

22. Ogbonna EC, Anderson HR, Beardslee PC, Bheemreddy P, Schmitz KR: Interactome Analysis Identifies MSMEI_3879 as a Substrate of Mycolicibacterium smegmatis ClpC1. Microbiology spectrum 2023, 11(4):e0454822.

23. Ogbonna EC, Anderson HR, Schmitz KR: Identification of Arginine Phosphorylation in Mycolicibacterium smegmatis. Microbiology spectrum 2022:e0204222.

24. Gopal P, Sarathy JP, Yee M, Ragunathan P, Shin J, Bhushan S, Zhu J, Akopian T, Kandror O, Lim TK et al: Pyrazinamide triggers degradation of its target aspartate decarboxylase. Nature communications 2020, 11(1):1661.

25. Ziemski M, Leodolter J, Taylor G, Kerschenmeyer A, Weber-Ban E: Genome-wide interaction screen for Mycobacterium tuberculosis ClpCP protease reveals toxin-antitoxin systems as a major substrate class. Febs j 2020.

26. Lunge A, Gupta R, Choudhary E, Agarwal N: The unfoldase ClpC1 of Mycobacterium tuberculosis regulates the expression of a distinct subset of proteins having intrinsically disordered termini. The Journal of biological chemistry 2020, 295(28):9455–9473.

27. Bachmair A, Finley D, Varshavsky A: In vivo half-life of a protein is a function of its amino-terminal residue. Science (New York, NY) 1986, 234(4773):179-186.

28. Tobias JW, Shrader TE, Rocap G, Varshavsky A: The N-end rule in bacteria. Science (New York, NY) 1991, 254(5036):1374-1377.

29. Varshavsky A: N-degron and C-degron pathways of protein degradation. Proceedings of the National Academy of Sciences of the United States of America 2019, 116(2):358–366.

30. Erbse A, Schmidt R, Bornemann T, Schneider-Mergener J, Mogk A, Zahn R, Dougan DA, Bukau B: ClpS is an essential component of the N-end rule pathway in Escherichia coli. Nature 2006, 439(7077):753-756.

31. Tasaki T, Sriram SM, Park KS, Kwon YT: The N-end rule pathway. Annual review of biochemistry 2012, 81:261–289.

32. Schuenemann VJ, Kralik SM, Albrecht R, Spall SK, Truscott KN, Dougan DA, Zeth K: Structural basis of N-end rule substrate recognition in Escherichia coli by the ClpAP adaptor protein ClpS. EMBO Rep 2009, 10(5):508–514.

33. Guo F, Esser L, Singh SK, Maurizi MR, Xia D: Crystal structure of the heterodimeric complex of the adaptor, ClpS, with the N-domain of the AAA+ chaperone, ClpA. The Journal of biological chemistry 2002, 277(48):46753–46762.

34. Stein BJ, Grant RA, Sauer RT, Baker TA: Structural Basis of an N-Degron Adaptor with More Stringent Specificity. Structure (London, England : 1993) 2016, 24(2):232-242.

35. Graciet E, Hu RG, Piatkov K, Rhee JH, Schwarz EM, Varshavsky A: Aminoacyl-transferases and the N-end rule pathway of prokaryotic/eukaryotic specificity in a human pathogen. Proceedings of the National Academy of Sciences of the United States of America 2006, 103(9):3078–3083.

36. Roman-Hernandez G, Grant RA, Sauer RT, Baker TA: Molecular basis of substrate selection by the N-end rule adaptor protein ClpS. Proceedings of the National Academy of Sciences of the United States of America 2009, 106(22):8888–8893.

37. Gao X, Yeom J, Groisman EA: The expanded specificity and physiological role of a widespread N-degron recognin. Proceedings of the National Academy of Sciences of the United States of America 2019, 116(37):18629–18637.

38. Stanne TM, Pojidaeva E, Andersson FI, Clarke AK: Distinctive types of ATP-dependent Clp proteases in cyanobacteria. The Journal of biological chemistry 2007, 282(19):14394–14402.

39. Nishimura K, Asakura Y, Friso G, Kim J, Oh SH, Rutschow H, Ponnala L, van Wijk KJ: ClpS1 is a conserved substrate selector for the chloroplast Clp protease system in Arabidopsis. Plant Cell 2013, 25(6):2276–2301.

40. Bouchnak I, van Wijk KJ: Structure, function, and substrates of Clp AAA+ protease systems in cyanobacteria, plastids, and apicoplasts: A comparative analysis. The Journal of biological chemistry 2021, 296:100338.

41. Tan JL, Ward L, Truscott KN, Dougan DA: The N-end rule adaptor protein ClpS from Plasmodium falciparum exhibits broad substrate specificity. FEBS Lett 2016, 590(19):3397–3406.

42. AhYoung AP, Koehl A, Vizcarra CL, Cascio D, Egea PF: Structure of a putative ClpS N-end rule adaptor protein from the malaria pathogen Plasmodium falciparum. Protein science : a publication of the Protein Society 2016, 25(3):689–701.

43. Wang KH, Oakes ES, Sauer RT, Baker TA: Tuning the strength of a bacterial N-end rule degradation signal. The Journal of biological chemistry 2008, 283(36):24600–24607.

44. Wang KH, Roman-Hernandez G, Grant RA, Sauer RT, Baker TA: The molecular basis of N-end rule recognition. Mol Cell 2008, 32(3):406–414.

45. Kunjapur AM, Stork DA, Kuru E, Vargas-Rodriguez O, Landon M, Söll D, Church GM: Engineering posttranslational proofreading to discriminate nonstandard amino acids. Proceedings of the National Academy of Sciences of the United States of America 2018, 115(3):619–624.

46. Alhuwaider AAH, Dougan DA: AAA+ Machines of Protein Destruction in Mycobacteria. Frontiers in molecular biosciences 2017, 4:49.

47. Guo C, Xiao Y, Bi F, Lin W, Wang H, Yao H, Lin D: Recombinant expression, biophysical and functional characterization of ClpS from Mycobacterium tuberculosis. Acta Biochim Biophys Sin (Shanghai) 2019, 51(11):1158–1167.

48. Finn RD, Clements J, Eddy SR: HMMER web server: interactive sequence similarity searching. Nucleic Acids Res 2011, 39(Web Server issue):W29–37.

49. Sievers F, Wilm A, Dineen D, Gibson TJ, Karplus K, Li W, Lopez R, McWilliam H, Remmert M, Soding J et al: Fast, scalable generation of high-quality protein multiple sequence alignments using Clustal Omega. Mol Syst Biol 2011, 7:539.

50. Wolf NM, Lee H, Zagal D, Nam JW, Oh DC, Lee H, Suh JW, Pauli GF, Cho S, Abad-Zapatero C: Structure of the N-terminal domain of ClpC1 in complex with the antituberculosis natural product ecumicin reveals unique binding interactions. *Acta crystallographica Section D*, Structural biology 2020, 76(Pt 5):458–471.

51. Jumper J, Evans R, Pritzel A, Green T, Figurnov M, Ronneberger O, Tunyasuvunakool K, Bates R, Žídek A, Potapenko A et al: Highly accurate protein structure prediction with AlphaFold. Nature 2021, 596(7873):583-589.

52. Jiang J, Schmitz KR: Bioinformatic identification of ClpI, a distinct class of Clp unfoldases in Actinomycetota. Frontiers in microbiology 2023, 14:1161764.

53. Waterhouse AM, Procter JB, Martin DM, Clamp M, Barton GJ: Jalview Version 2--a multiple sequence alignment editor and analysis workbench. Bioinformatics (Oxford, England) 2009, 25(9):1189–1191.

54. Crooks GE, Hon G, Chandonia JM, Brenner SE: WebLogo: a sequence logo generator. Genome research 2004, 14(6):1188–1190.

55. Ashkenazy H, Abadi S, Martz E, Chay O, Mayrose I, Pupko T, Ben-Tal N: ConSurf 2016: an improved methodology to estimate and visualize evolutionary conservation in macromolecules. Nucleic Acids Res 2016, 44(W1):W344–350.

56. Glaser F, Pupko T, Paz I, Bell RE, Bechor-Shental D, Martz E, Ben-Tal N: ConSurf: identification of functional regions in proteins by surface-mapping of phylogenetic information. *Bioinformatics (Oxford*, England) 2003, 19(1):163–164.

57. Gibson DG, Young L, Chuang RY, Venter JC, Hutchison CA, 3rd, Smith HO: Enzymatic assembly of DNA molecules up to several hundred kilobases. Nature methods 2009, 6(5):343–345.

58. Hershfield V, Boyer HW, Yanofsky C, Lovett MA, Helinski DR: Plasmid ColEl as a molecular vehicle for cloning and amplification of DNA. Proceedings of the National Academy of Sciences of the United States of America 1974, 71(9):3455–3459.

59. Guo XV, Monteleone M, Klotzsche M, Kamionka A, Hillen W, Braunstein M, Ehrt S, Schnappinger D: Silencing Mycobacterium smegmatis by using tetracycline repressors. J Bacteriol 2007, 189(13):4614–4623.

60. Kaps I, Ehrt S, Seeber S, Schnappinger D, Martin C, Riley LW, Niederweis M: Energy transfer between fluorescent proteins using a co-expression system in Mycobacterium smegmatis. Gene 2001, 278(1-2):115–124.

61. Ehrt S, Guo XV, Hickey CM, Ryou M, Monteleone M, Riley LW, Schnappinger D: Controlling gene expression in mycobacteria with anhydrotetracycline and Tet repressor. Nucleic Acids Res 2005, 33(2):e21.

62. Lau YK, Baytshtok V, Howard TA, Fiala BM, Johnson JM, Carter LP, Baker D, Lima CD, Bahl CD: Discovery and engineering of enhanced SUMO protease enzymes. The Journal of biological chemistry 2018, 293(34):13224–13233.

63. Zhu Y, Mao C, Ge X, Wang Z, Lu P, Zhang Y, Chen S, Hu Y: Characterization of a Minimal Type of Promoter Containing the −10 Element and a Guanine at the −14 or −13 Position in Mycobacteria. J Bacteriol 2017, 199(21).

64. Rock JM, Hopkins FF, Chavez A, Diallo M, Chase MR, Gerrick ER, Pritchard JR, Church GM, Rubin EJ, Sassetti CM et al: Programmable transcriptional repression in mycobacteria using an orthogonal CRISPR interference platform. Nature microbiology 2017, 2:16274.

65. Malakhov MP, Mattern MR, Malakhova OA, Drinker M, Weeks SD, Butt TR: SUMO fusions and SUMO-specific protease for efficient expression and purification of proteins. Journal of structural and functional genomics 2004, 5(1-2):75–86.

66. Otwinowski Z, Minor W: Processing of X-ray Diffraction Data Collected in Oscillation Mode. In: Macromolecular Crystallography. Edited by Carter Jr. CW, Sweet RM, vol. 276. New York: Academic Press; 1997: 307–326.

67. McCoy AJ, Grosse-Kunstleve RW, Adams PD, Winn MD, Storoni LC, Read RJ: Phaser crystallographic software. J Appl Crystallogr 2007, 40(Pt 4):658–674.

68. Roman-Hernandez G, Hou JY, Grant RA, Sauer RT, Baker TA: The ClpS adaptor mediates staged delivery of N-end rule substrates to the AAA+ ClpAP protease. Mol Cell 2011, 43(2):217–228.

69. Emsley P, Cowtan K: Coot: model-building tools for molecular graphics. *Acta crystallographica Section D*, Biological crystallography 2004, 60(Pt 12 Pt 1):2126–2132.

70. Adams PD, Afonine PV, Bunkoczi G, Chen VB, Davis IW, Echols N, Headd JJ, Hung LW, Kapral GJ, Grosse-Kunstleve RW et al: PHENIX: a comprehensive Python-based system for macromolecular structure solution. *Acta crystallographica Section D*, Biological crystallography 2010, 66(Pt 2):213–221.

71. Hou JY, Sauer RT, Baker TA: Distinct structural elements of the adaptor ClpS are required for regulating degradation by ClpAP. Nature structural & molecular biology 2008, 15(3):288–294.

72. Rivera-Rivera I, Román-Hernández G, Sauer RT, Baker TA: Remodeling of a delivery complex allows ClpS-mediated degradation of N-degron substrates. Proceedings of the National Academy of Sciences of the United States of America 2014, 111(37):E3853–3859.

73. Zeth K, Ravelli RB, Paal K, Cusack S, Bukau B, Dougan DA: Structural analysis of the adaptor protein ClpS in complex with the N-terminal domain of ClpA. Nat Struct Biol 2002, 9(12):906–911.

74. Marsee JD, Ridings A, Yu T, Miller JM: Mycobacterium tuberculosis ClpC1 N-Terminal Domain Is Dispensable for Adaptor Protein-Dependent Allosteric Regulation. Int J Mol Sci 2018, 19(11).

75. Varshavsky A: The N-end rule pathway and regulation by proteolysis. Protein science : a publication of the Protein Society 2011, 20(8):1298–1345.

76. H. Ashkenazy, E. Erez, E. Martz, T. Pupko, Ben-Tal N: ConSurf 2010: calculating evolutionary conservation in sequence and structure of proteins and nucleic acids. Nucleic Acids Res 2010, 38(Web Server issue):W529–533.

77. Li Y, Corro JH, Palmer CD, Ojha AK: Progression from remodeling to hibernation of ribosomes in zinc-starved mycobacteria. Proceedings of the National Academy of Sciences of the United States of America 2020, 117(32):19528–19537.

78. Ichetovkin IE, Abramochkin G, Shrader TE: Substrate recognition by the leucyl/phenylalanyl-tRNA-protein transferase. Conservation within the enzyme family and localization to the trypsin-resistant domain. The Journal of biological chemistry 1997, 272(52):33009–33014.

79. Leibowitz MJ, Soffer RL: A soluble enzyme from Escherichia coli which catalyzes the transfer of leucine and phenylalanine from tRNA to acceptor proteins. Biochemical and biophysical research communications 1969, 36(1):47–53.

80. Olaleye O, Raghunand TR, Bhat S, He J, Tyagi S, Lamichhane G, Gu P, Zhou J, Zhang Y, Grosset J et al: Methionine aminopeptidases from Mycobacterium tuberculosis as novel antimycobacterial targets. Chem Biol 2010, 17(1):86–97.

81. Yan MY, Li SS, Ding XY, Guo XP, Jin Q, Sun YC: A CRISPR-Assisted Nonhomologous End-Joining Strategy for Efficient Genome Editing in Mycobacterium tuberculosis. mBio 2020, 11(1).

82. Yan MY, Yan HQ, Ren GX, Zhao JP, Guo XP, Sun YC: CRISPR-Cas12a-Assisted Recombineering in Bacteria. Applied and environmental microbiology 2017, 83(17).

83. Kim S, Fei X, Sauer RT, Baker TA: AAA+ protease-adaptor structures reveal altered conformations and ring specialization. Nature structural & molecular biology 2022, 29(11):1068–1079.

84. Xiao Q, Zhang F, Nacev BA, Liu JO, Pei D: Protein N-terminal processing: substrate specificity of Escherichia coli and human methionine aminopeptidases. Biochemistry 2010, 49(26):5588–5599.

85. Hirel PH, Schmitter MJ, Dessen P, Fayat G, Blanquet S: Extent of N-terminal methionine excision from Escherichia coli proteins is governed by the side-chain length of the penultimate amino acid. Proceedings of the National Academy of Sciences of the United States of America 1989, 86(21):8247–8251.

86. Ben-Bassat A, Bauer K, Chang SY, Myambo K, Boosman A, Chang S: Processing of the initiation methionine from proteins: properties of the Escherichia coli methionine aminopeptidase and its gene structure. J Bacteriol 1987, 169(2):751–757.

87. Zhang X, Chen S, Hu Z, Zhang L, Wang H: Expression and characterization of two functional methionine aminopeptidases from Mycobacterium tuberculosis H37Rv. Current microbiology 2009, 59(5):520–525.

