## Supplementary Figures S1 - S6 for "ClpS directs degradation of primary N-end rule substrates in *Mycolicibacterium smegmatis*"

#### Figure S1

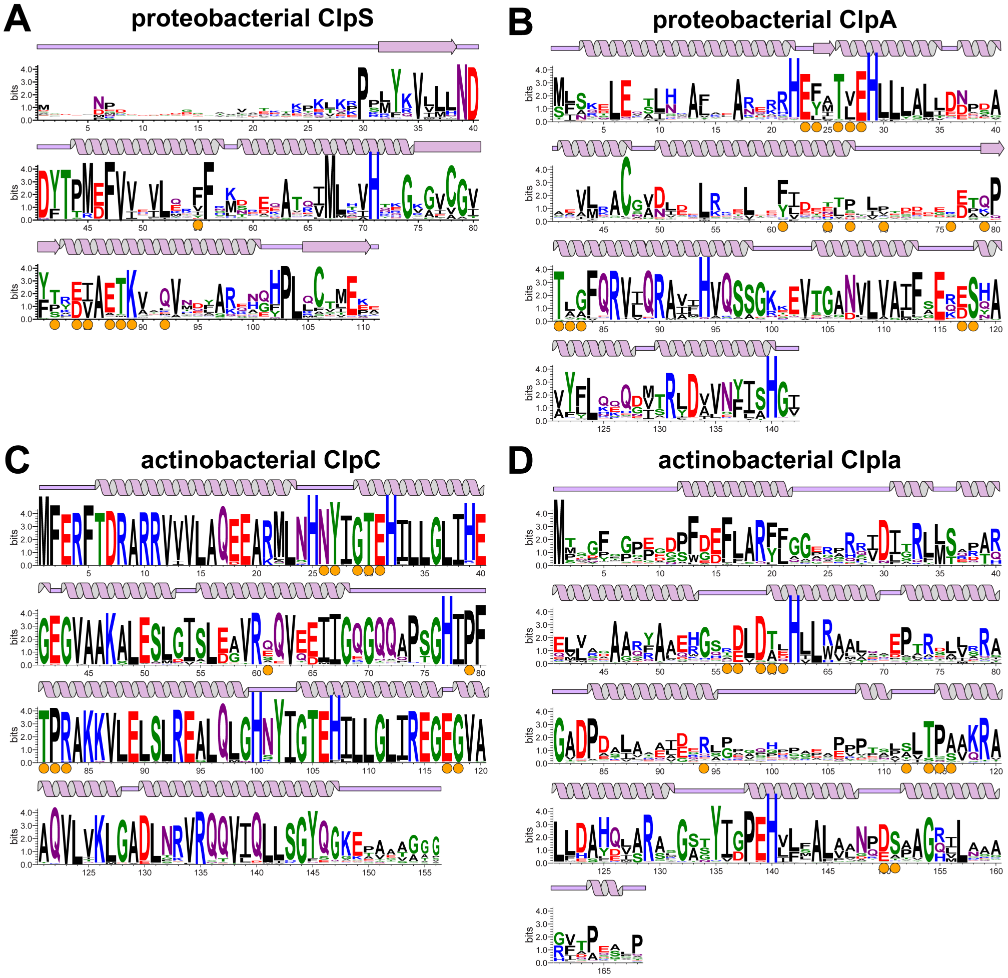


**Figure S1. Sequence conservation among ClpS and unfoldase orthologs**. The search tool HMMER [48] was used to compile sequence alignments of **A**) proteobacterial ClpS, **B**) proteobacterial ClpA, **C**) actinobacterial ClpC or **D**) actinobacterial ClpIa, which were used to construct the resulting sequence logos using WebLogo [54]. Plots of approximate secondary structure are derived from crystal structures of *E. coli* ClpS and the ClpA NTD (PDB: 1MBU) [33], the *M. tuberculosis* ClpC1 NTD (PDB: 6PBA) [50], or an AlphaFold2 [51] model of *Streptomyces coelicoor* ClpIa (Uniprot: O69936) [52].

#### Figure S2

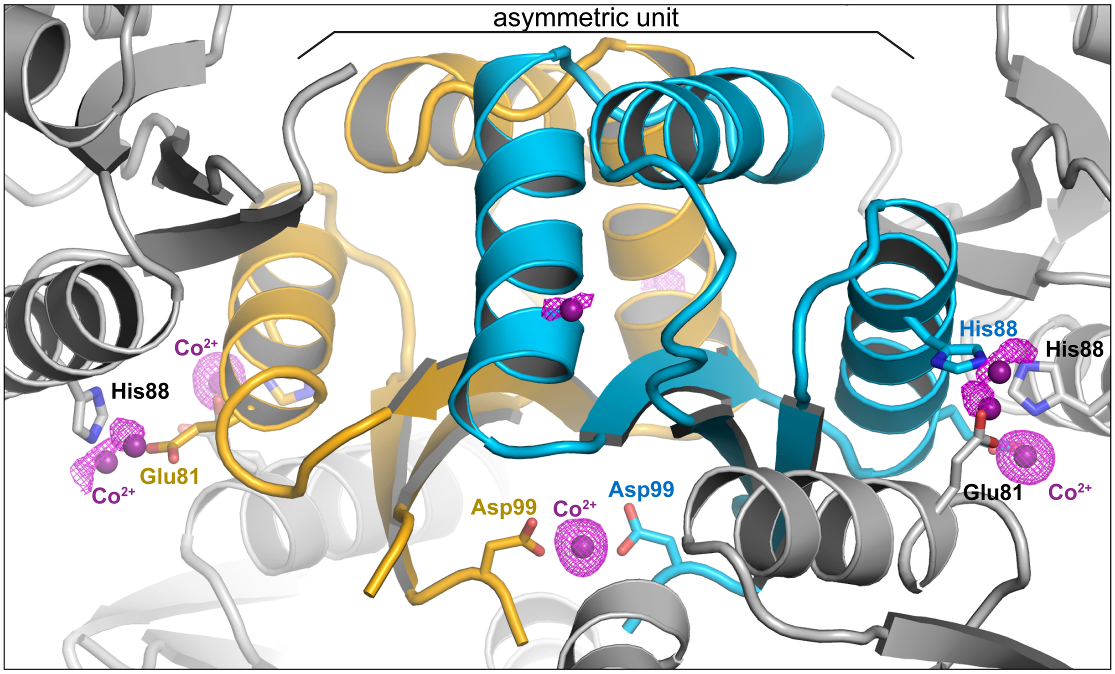


**Figure S2. Anomalous map of *M. smegmatis* ClpS showing bound cobalt ions**. An anomolous electron density map (purple; contour 3.5 σ) is shown over a cartoon of the *M. smegmatis* ClpS$^{core}$ crystal structure, with asymmetric unit chains in gold and cyan, symmetry related chains colored gray, and $Co^{2+}$ ions shown as purple spheres. Key residues interacting with cobalt ions are labeled and shown as sticks.

#### Figure S3

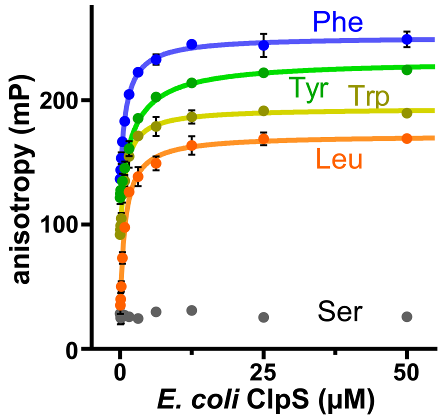


**Figure S3. Binding of *E. coli* ClpS to N-end rule peptides**. Fluorescence anisotropy of 100 nM TAMRA-labeled peptides bearing the indicated N-terminal amino acid was monitored across increasing concentrations of *E. coli* ClpS. Data were fit to a single site binding equation. ClpS bound peptides with N-term Leu with $K_D$ = 0.88 ± 0.07 µM, Tyr with $K_D$ = 2.5 ± 0.2 µM, Trp with $K_D$ = 0.95 ± 0.08 µM, Phe with $K_D$ = 1.0 ± 0.08 µM. No binding was observed to a peptide with N-term Ser.

### Figure S4

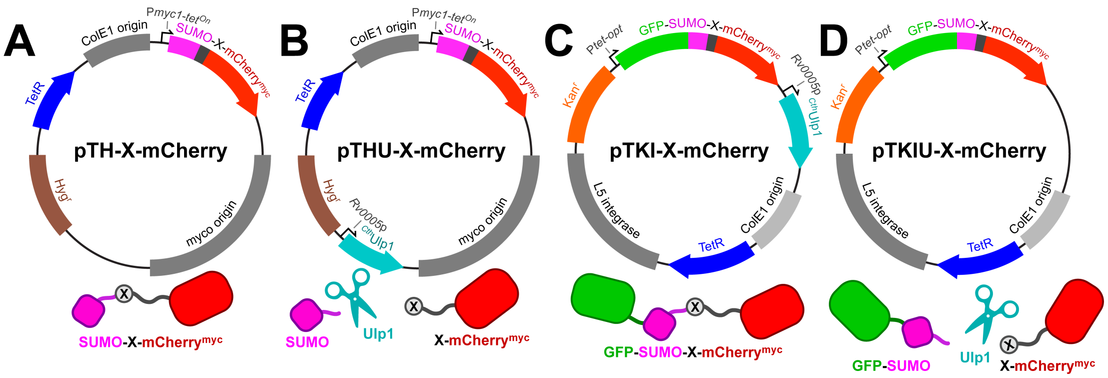


**Figure S4. Plasmid maps of mycobacterial expression constructs**. Non-integrative mycobacterial shuttle plasmids for the expression of SUMO-X-mCherry$^{myc}$ **A**) without and **B**) with co-expression of Ulp1. Integrative mycobacterial shuttle plasmids for the expression of GFP-SUMO-X-mCherry$^{myc}$ **C**) without and **D**) with co-expression of Ulp1.

#### Figure S5

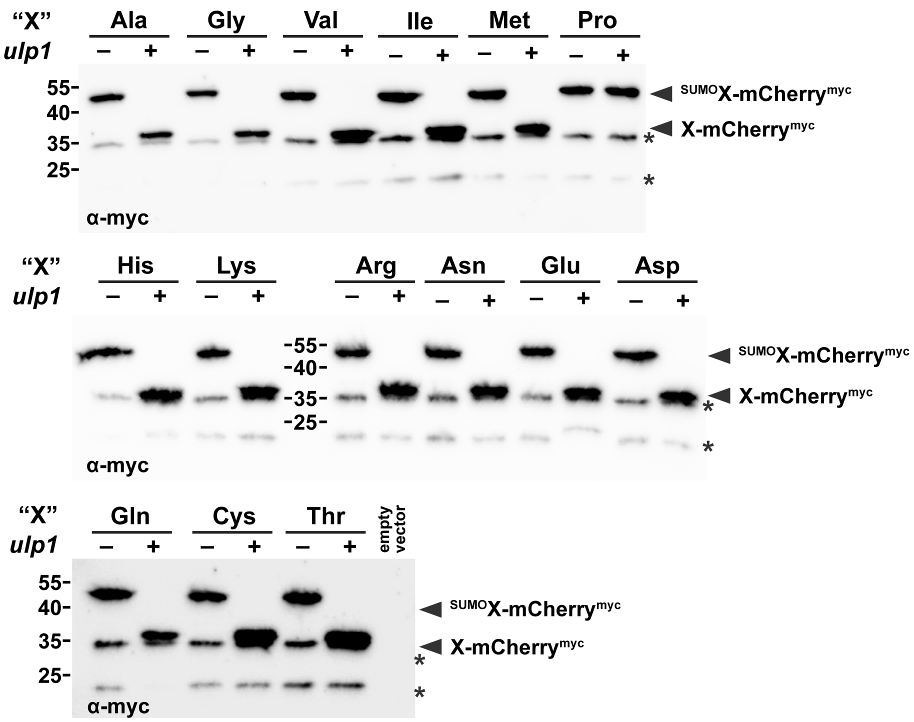


**Figure S5. Expression of SUMO-X-mCherry$^{myc}$ constructs**. Immunoblots of lysates from cells expressing the indicated SUMO-X-mCherry$^{myc}$ constructs, with or without co-expressed Ulp1, were probed with an α-myc antibody. With the exception of SUMO-Pro-mCherry$^{myc}$, all constructs migrate at as smaller species in the presence of Ulp1, consistent with removal of the shielding SUMO. Asterisks indicates truncation products.

#### Figure S6

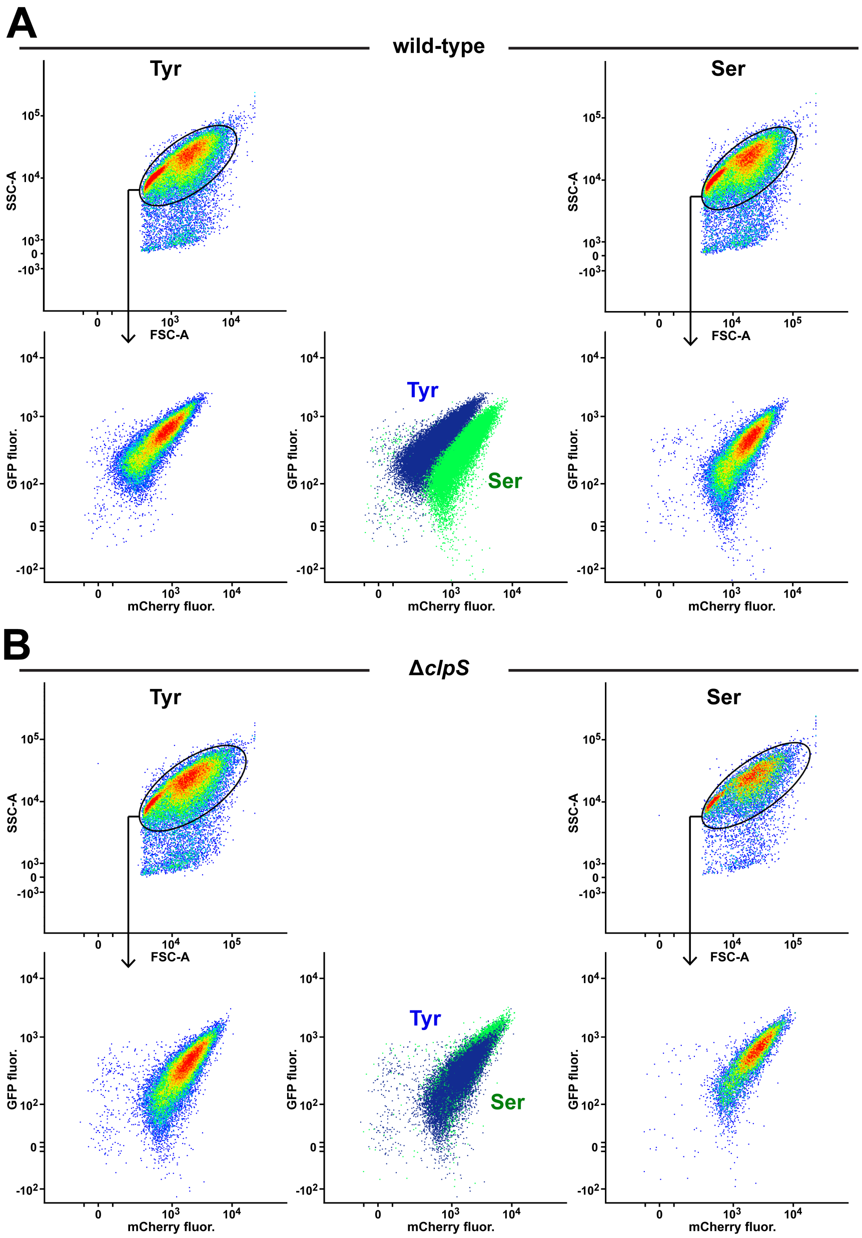


**Figure S6. Flow cytometry analysis of GFP-SUMO-X-mCherry$^{myc}$.** Light scattering and fluorescence of GFP-SUMO-Tyr-mCherry$^{myc}$ (left) and GFP-SUMO-Ser-mCherry$^{myc}$ (right) constructs with co-expressed Ulp1 in **A**) wild-type and **B**) Δ*clpS M. smegmatis* was analyzed by flow cytometry. Events were filtered by elliptical gates on the basis of side (SSC-A) and forward scattering (FSC-A) that captured ~90% of events.
